# High-resolution imaging reveals a cascade of interconnected cellular bioeffects differentiating the long-term fates of sonoporated cells

**DOI:** 10.1101/2025.05.08.652390

**Authors:** Jonathan LF Lee, Jae Song, Rumelo Amor, Jonathon Bolton, Andrew Thompson, Jürgen Götz, Pranesh Padmanabhan

**Author notes:** **Correspondence:** Pranesh Padmanabhan; Jürgen Götz.

## Abstract

Low-intensity ultrasound combined with microbubbles is a promising, non-invasive treatment strategy for enhancing vascular permeability and targeted intracellular drug or gene delivery. After ultrasound insonification, cells can undergo reversible sonoporation, involving adaptation and recovery, or irreversible sonoporation, marked by a loss of cell viability. To design effective sonoporation-based therapeutic delivery strategies, it is therefore critical to identify the biological responses that determine these distinct cell fates. Here, we developed a custom-built high-resolution multicolour imaging device and applied a single pulse of ultrasound (267 kHz frequency, 20 μs duration, and ∼110 kPa peak-negative pressure) to trigger targeted microbubble-mediated reversible and irreversible sonoporation events within a confluent monolayer of cultured Madin-Darby canine kidney (MDCK) II cells. We found that intracellular calcium levels rose rapidly and peaked within 10 s in both types of sonoporated cells, although the levels declined differently. In reversibly sonoporated cells, these levels gradually returned to baseline, whereas in irreversibly sonoporated cells, they dropped rapidly, falling well below baseline within 7.9 ± 3.0 min (mean ± s.d.). Using single-vesicle imaging, we further found that vesicles containing the tight junction protein claudin-5 remained mobile with subtly reduced movement in reversibly sonoporated cells, whereas they almost stalled in irreversibly sonoporated cells. The underlying microtubule network was partially disrupted in the reversibly sonoporated cells, recovering fully within 3.2 ± 2.9 min (mean ± s.d.). In contrast, in irreversibly sonoporated cells, the entire microtubule network collapsed within 4.0 ± 2.4 min (mean ± s.d.). Whilst in reversibly sonoporated cells, the uptake of the model drug propidium iodide was mild-to-moderate, without drastic cell size changes up to about 1 h post-sonication, irreversibly sonoporated cells presented with substantially higher propidium iodide uptake and shrunk completely within 43.2 ± 10.5 min (mean ± s.d.). Together, our study identified distinct spatiotemporal sequences of interconnected biological responses underlying different fates of sonoporated cells, providing a framework for identifying processes that could be manipulated for safe and effective sonoporation-based drug delivery.

## INTRODUCTION

Sonoporation utilises ultrasound-mediated membrane permeabilisation as a means by which drugs or genes can be delivered to specific targets. This promising strategy is currently under preclinical and clinical investigation across diverse medical fields, including neurology, cardiology, and oncology [1–5]. Upon ultrasound exposure, due to pressure changes induced by ultrasound waves, exogenously added microbubbles expand and contract rapidly [6,7], thereby exerting various mechanical and biophysical forces on cells [8]. This can lead to membrane pore formation, termed sonoporation [9], enabling intracellular delivery of therapeutics [8,10–12]. Microbubble stimulation can also increase vesicle-mediated transcellular transport of therapeutics [13,14], as well as enhance vascular permeability by opening the endothelial cell-cell contacts that form the blood-brain barrier [15,16]. Of the many microbubble-mediated effects, it has been shown that sonoporation triggers a complex cascade of molecular and cellular responses, termed sonoporation-induced bioeffects, which alter cellular functions at multiple spatial and temporal scales [1,17–19]. This collectively determines whether cells undergo reversible sonoporation, involving adaptation and recovery, or irreversible sonoporation, characterised by cell death [17]. In specific clinical settings, such as delivering therapeutics to cancer cells [20–22], a higher occurrence of irreversible sonoporation may offer an additional therapeutic advantage by reducing the tumour burden. In contrast, in other settings, such as targeting the brain via the opening of the blood-brain barrier [23–27], it may be critical to minimise irreversible sonoporation to ensure biosafety. Therefore, identifying the underlying cellular bioeffects and their spatiotemporal interdependence that determine the fates of sonoporated cells is crucial for the rational design of sonoporation-based treatment strategies that effectively balance the trade-off between delivery effectiveness and safety.

Optical microscopy studies have provided significant insights into the complex dynamic behaviour of microbubbles under different combinations of ultrasound parameters [28–30], their interactions with cells [28–31], and subsequent membrane pore formation [10,28–31]. They have also characterised downstream bioeffects, including transient changes in intracellular calcium levels [32–38], alterations in reactive oxygen species levels [35,39], endoplasmic reticulum stress [40], and cytoskeletal rearrangements [35,41–44]. These observations have been instrumental in linking some of the observed molecular and cellular responses to cell recovery processes, such as membrane resealing [10,45,46], and cell morphological changes that promote drug delivery [47], such as intercellular gap formation [29,35,43,48] and transcellular perforation [43,48,49]. More recently, an optical imaging-based study showed that manipulating intracellular calcium levels could positively influence long-term functions, such as cell proliferation, of reversibly sonoporated cells [38]. However, despite these conceptual advances, a detailed understanding of the cascade of bioeffects differentiating reversible and irreversible sonoporation is still lacking.

In this study, we developed and validated a custom-built system combining a 267 kHz single-element ultrasound transducer with an inverted laser scanning confocal microscope with high-resolution imaging capabilities. Using this setup, we stimulated targeted microbubbles with a single ultrasound pulse, triggered reversible and irreversible sonoporation events within a monolayer of cultured Madin-Darby canine kidney (MDCK) II cells, and quantified the downstream bioeffects in the two types of sonoporated cells for up to ∼1 h. This approach enabled us to identify spatially and temporally interconnected changes in intracellular calcium response, vesicle movement, microtubule network dynamics, propidium iodide (PI) uptake, and cell morphodynamics that differentiate the fates of sonoporated cells.

## RESULTS

### Device development for real-time imaging of bioeffects triggered by ultrasound-induced sonoporation

To quantify sonoporation-induced molecular and cellular changes in live cells, we first characterised a low-frequency ultrasound transducer in the free-field (Fig. 1), then developed a custom-built device coupling this transducer with an inverted high-resolution confocal microscope (Fig. 2), and generated and characterised targeted microbubbles (Fig. 3).

**Figure 1.**
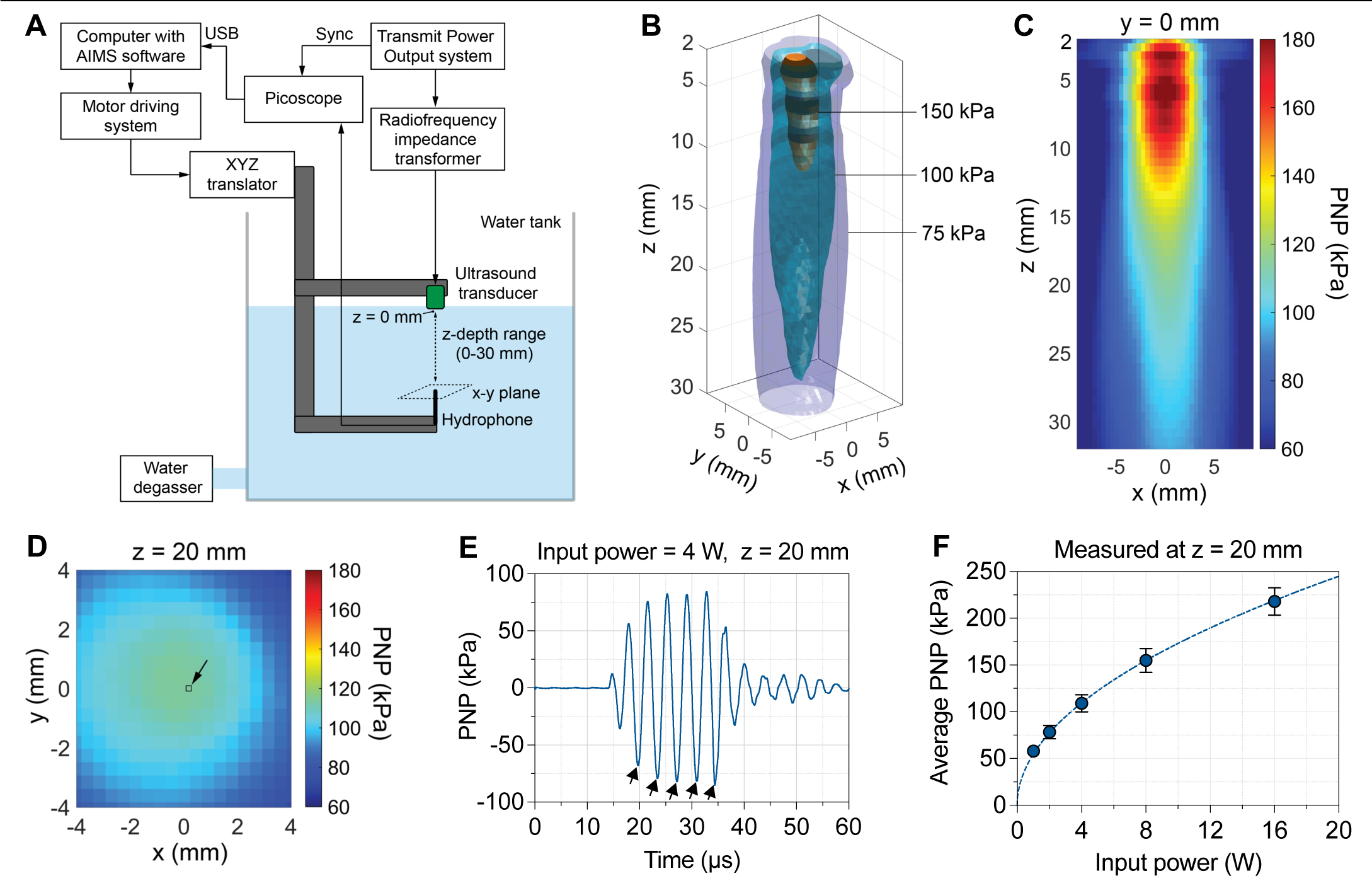
Characterisation of a single-element 267 kHz ultrasound transducer in a free-field tank system. **(A)** Schematic of the setup consisting of a transducer scanning apparatus in a scanning tank (Onda Corp., USA) filled with degassed water to characterise the focus and acoustic field generated by a single-element transducer (Fraunhofer, IBTM, Germany). Ultrasound waveforms received by the hydrophone (HNR-1000, Onda Corp., USA) were recorded by a computer-controlled oscilloscope in synchronisation with ultrasound excitation. The unfocussed, single-element transducer was driven by a transmitter power output (TPO) system (TPO^TM^, Sonic Concepts Inc., USA) and a radiofrequency impedance transformer to generate ultrasound pulses. **(B)** 3D contours representing the acoustic field at PNP levels of 75, 100 and 150 kPa were obtained by scanning along the x, y and z axes. **(C)** 2D map of PNP along the x and z axes at y = 0. **(D)** 2D map of PNP along the x and y axes at a z-depth of 20 mm away from the transducer surface. The black square represents the field of view (211.5 x 211.5 µm) in our imaging setup. **(E)** The sinusoidal waveform generated by a single, 20 µs short ultrasound pulse at a z-depth of 20 mm and with an input frequency of 267 kHz and a peak input power of 4 W. The black arrows indicate the five PNPs used to calculate the average PNP displayed in panel F. **(F)** The average PNP as a function of input power to the TPO system measured at a z-depth of 20 mm. The circles and error bars are the mean and s.d. of PNPs corresponding to the five cycles shown in E. The dashed line is the best fit of the equation 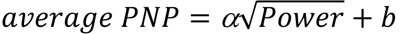 to the data.

**Figure 2.**
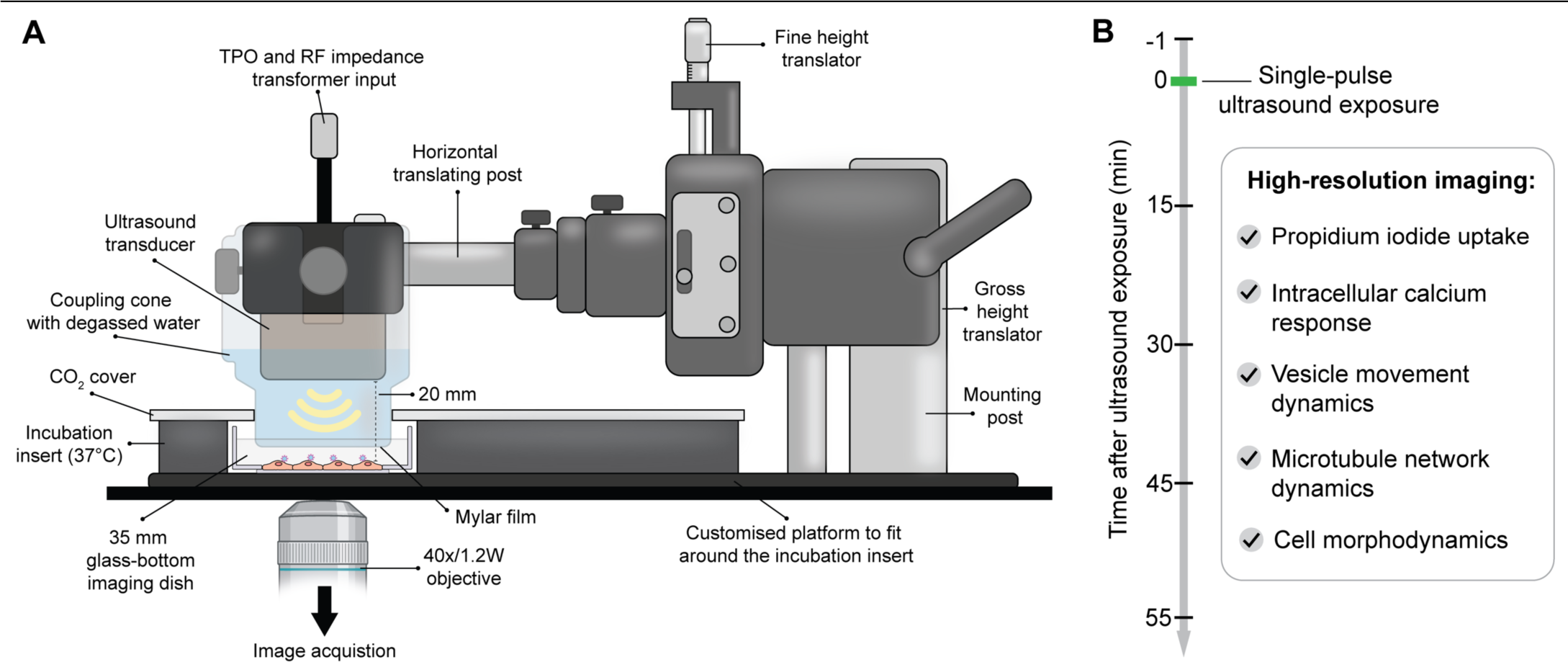
Schematic of the custom-built device used to sonicate cells and perform live-cell imaging. **(A)** The device combines the Zeiss LSM980 laser scanning confocal microscope with non-linear optics and an Airyscan 2 detector, and a single-element ultrasound transducer (Fraunhofer IBMT) immersed in a degassed water-containing coupling cone. The bottom membrane of the coupling cone interfaces with the cell culture in the imaging dish. The distance between the transducer and the cells was set at 20 mm, where the PNP variation along the xy-plane is minimal (Fig. 1D). **(B)** The custom-built platform was used to expose cells to ultrasound and quantify several key downstream bioeffects.

**Figure 3.**
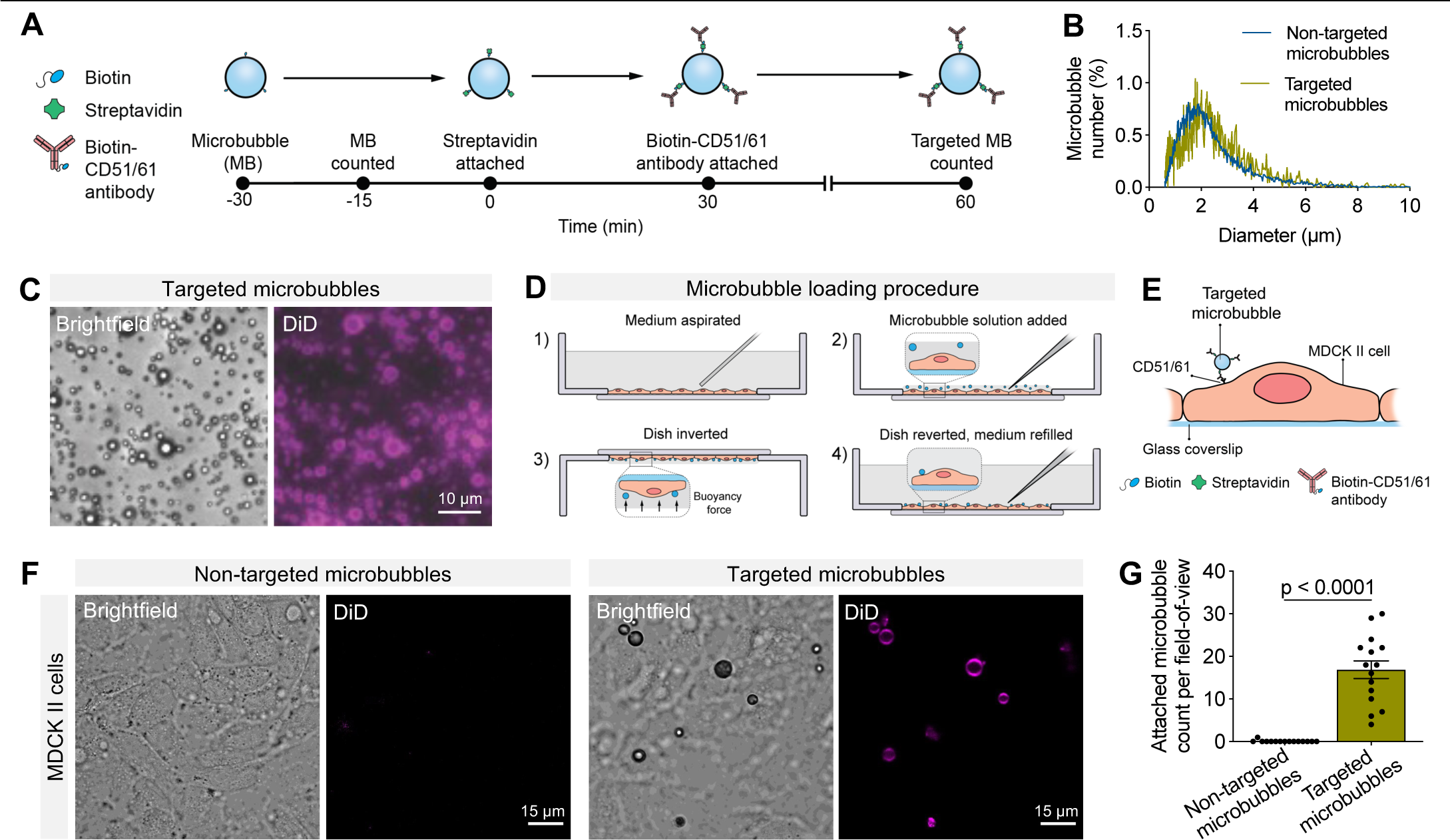
Generation and characterisation of targeted microbubbles and their attachment to MDCK II cells. **(A)** Timeline of microbubble functionalisation with an anti-CD51/61 antibody through biotin-streptavidin bridging. **(B)** Size distribution of targeted and non-targeted microbubbles determined in solution. **(C)** Brightfield and fluorescent images of targeted microbubbles labelled with the lipid dye DiD in glycerol. **(D)** Schematic of the loading procedure used to attach targeted microbubbles to the apical surface of MDCK II cells cultured in a 35 mm glass-bottom dish. **(E)** Schematic of a simplified *in situ* scenario in which a targeted microbubble is bound to the apical surface of an MDCK II cell. **(F)** Representative images of non-targeted and targeted microbubbles, visualised by brightfield and DiD fluorescence, near the confluent MDCK II cell monolayers. Non-targeted microbubbles did not attach to cells. **(G)** Number of non-targeted or targeted microbubbles attached to MDCK II cells per field of view (211.5 x 211.5 µm). Error bars, s.e.m. n = 15 fields of view for each condition from 2 independent experiments.

As ultrasound treatment in the ∼220-650 kHz frequency range has recently entered clinical trials [50,51] and microbubbles are also excited in the kHz frequency range [52–54], we first sought to characterise a bespoke unfocussed, single-element, circular transducer (Fraunhofer IBMT) measuring 18 mm in diameter and operating at a central frequency of 267 kHz in a degassed water tank (Fig. 1A; Methods). Using an Onda HNR-1000 hydrophone, we measured the 3-dimensional (3D) acoustic pressure field generated by the transducer and computed 3D contours corresponding to the peak-negative pressure (PNP) of 75, 100 and 150 kPa (Fig. 1B). As expected, the 2D and 3D maps revealed that the PNP was maximal near the transducer surface and decreased along the x, y and z axes (Fig. 1B). Notably, the variation in the PNP levels in the xy-plane decreased with z-depth. We set the z-depth at 20 mm for subsequent imaging work (Fig. 1C), as the PNP variation along the xy-plane across the imaging field of view of size 211.5 x 211.5 µm was less than 1% (Fig. 1D). At a z-depth of 20 mm for a peak input power of 4 W, a single 20 µs ultrasound pulse generated a pressure waveform comprising five cycles, with the average PNP equalling 109.1 ± 9.1 kPa (mean ± s.d.) (Fig. 1E). Such a short pulse has routinely been used to trigger single-site sonoporation [10,29,32–34,36–39,41–46,48,49,55]. The average PNP monotonically increased with the input power, and the equation 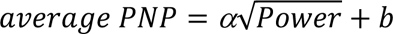, where *α* and *b* are constants, provided a good fit to the data (Fig. 1F), indicating a square root PNP-power relationship.

We next assembled a custom-built device consisting of the above-described ultrasound transducer immersed in a coupling cone filled with degassed water and paired it with an inverted laser-scanning high-resolution confocal microscope. The bottom membrane of the coupling cone interfaced with the culture medium in the imaging dish with controlled temperature, CO_2_ and humidity levels (Fig. 2, Supplementary Fig. S1). This setup allowed us to deliver ultrasound from above while imaging molecular and cellular processes from below for up to 1 h post-sonication and investigate the diversity of sonoporation-induced bioeffects across cells at a fixed ultrasound frequency and acoustic pressure.

Having established a system to monitor cellular responses to ultrasound, we next conjugated microbubbles with a ligand for a cell surface receptor to target them to the apical cell surface for examining sonoporation-dependent downstream bioeffects [32,34,49]. We chose MDCK II cells as a model cellular system, given their ability to form a functional monolayer [56] and their previous use in ultrasound-mediated drug delivery studies [57,58]. As previous proteomic studies have reported CD51/61 expression in MDCK cells [59,60], we generated targeted microbubbles by functionalising their lipid shell with a biotin-streptavidin bridge coupled to a biotinylated anti-CD51/61 antibody (Fig. 3A). We incorporated a fluorescent lipid dye DiD to locate the microbubbles in the far-red channel (639 nm) and found that CD51/61 antibody conjugation into the bubble shell did not alter the microbubble size distribution in saline solution (Fig. 3B). Using images captured in both brightfield and the far-red channel, we confirmed the spherical structure and polydispersity of the microbubbles, which were immobilised in glycerol to avoid motion artefacts during imaging (Fig. 3C). We next administered a small aliquot of targeted microbubbles (using non-targeted microbubbles as a control) to the MDCK II cell monolayer cultured in 35 mm glass-bottom imaging dishes, inverted the dishes to facilitate microbubble attachment to the cell surface by flotation, after which we reverted the dish to wash away any excess or unattached microbubbles (Fig. 3E-F). On average, we detected 16.9 ± 8.0 (mean ± s.d.) targeted microbubbles attached to cells per field of view, whereas we hardly detected any non-targeted microbubbles attached to cells, yielding an average of 0.1 ± 0.3 (mean ± s.d.) per field of view (Fig. 3F-G). These observations demonstrated that targeted microbubbles robustly attach to MDCK II cells. Next, we examined the responses of cells with and without targeted microbubbles attached to ultrasound exposure.

### Distinct dynamics of intracellular calcium responses in reversibly and irreversibly sonoporated cells

Previous imaging studies have observed transient elevations of intracellular calcium levels following sonoporation-induced membrane perforation [32–38], but the long-term dynamics in sonoporated cells remain poorly understood. To address this question and validate our experimental setup, we generated an MDCK II cell line (GCaMP6f-MDCK II) that stably expresses GCaMP6f, a genetically encoded reporter for cytosolic calcium levels [61]. After adding targeted microbubbles to a confluent monolayer of GCaMP6f-MDCK II cells, we identified cells with and without microbubbles attached to their surfaces via the DiD signal (639 nm, far-red channel). Next, we performed time-lapse imaging of intracellular calcium responses (488 nm, green channel) and sonoporation via the uptake of the model drug PI (561 nm, red channel). We recorded baseline values for 1 min before sonication and sonoporation-dependent changes in calcium and PI levels for 55 min. The sonication consisted of a short ultrasound pulse of 20 μs duration at a frequency of 267 kHz and a PNP of ∼110 kPa. This was sufficient to elicit sonoporation in many spatially distant cells within the field of view.

In cells without microbubbles attached, we observed neither PI uptake nor changes in intracellular calcium levels in response to sonication (Fig. 4A; Video S1). We refer to this control group as non-sonoporated cells. In ∼46 % of the cells with targeted microbubbles, we observed a rapid PI uptake accompanied by a transient increase in intracellular calcium levels (Fig. 4B-C; Videos S2 and S3). The rapid PI uptake started at a confined region, and from there, the PI signal gradually spread throughout the entire cell within a couple of minutes. The site of PI entry closely matched the location of the microbubble attachment (Fig. 4B-C; Supplementary Fig. S2), indicating that the dynamics of PI uptake is a consequence of ultrasound-induced sonoporation. While we observed intercellular calcium waves, where the calcium signals propagated to neighbouring non-sonoporated cells, the increase in PI levels was confined solely to the sonoporated cells (Fig. 4B-C; Supplementary Fig. S3). The calcium elevation in the neighbouring cells was transient and returned to baseline within 10 min (Supplementary Fig. S3). Crucially, based on the dynamics of cell morphological changes, we identified at least two distinct subpopulations of sonoporated cells (Fig. 4B-C), which we refer to as reversibly and irreversibly sonoporated cells for the following reasons. In reversibly sonoporated cells, we observed a significant increase in PI levels that plateaued within 5 min (Fig. 4D). The levels in these cells were higher than in non-sonoporated cells but substantially lower than those in irreversibly sonoporated cells (Fig. 4E). Concomitantly, intracellular calcium levels rose rapidly, reaching peak intensities within ∼10 s, and gradually returned close to baseline levels within ∼40 min (Fig. 4F). Moreover, the intracellular calcium elevation, as assessed using the area under the relative GCaMP6f intensity curve (AUC), strongly correlated with PI levels, indicating that sonoporation-induced membrane perforation, which causes PI uptake, and the calcium response are interlinked (Fig. 4F inset). Importantly, considering that the GCaMP6f signal remained detectable throughout the remainder of the imaging (i.e., for 55 min), and that cytosolic calcium is a marker of cell viability, these cells thus underwent reversible sonoporation, presumably without compromised viability.

**Figure 4.**
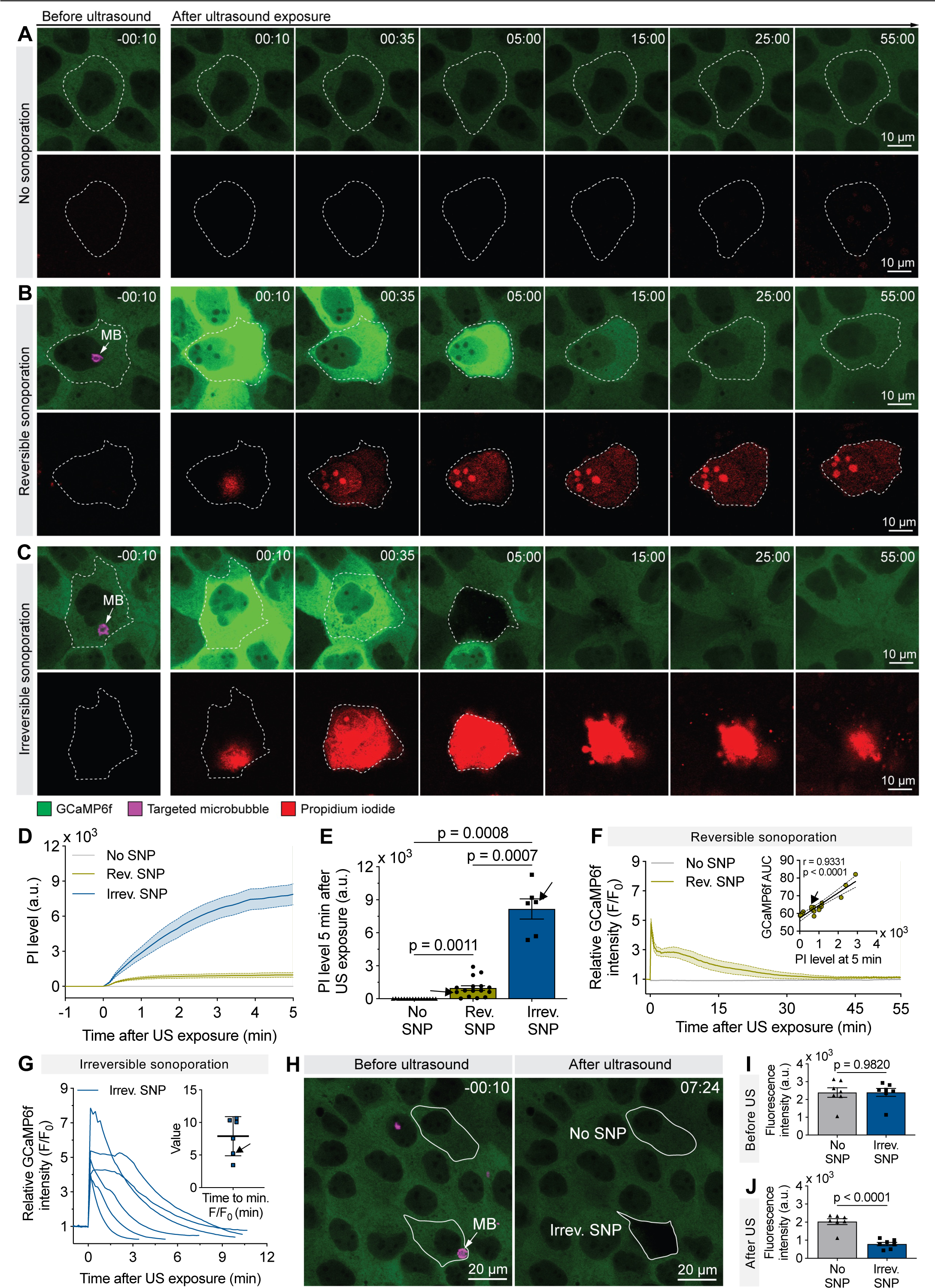
Intracellular calcium responses are distinct in reversibly and irreversibly sonoporated cells. **(A-C)** Representative time-lapse images of GCaMP6f fluorescence intensity representing intracellular calcium levels and the corresponding PI levels before and after sonication in non-(A), reversibly (B) and irreversibly (C) sonoporated cells. Dashed white lines demarcate cellular outlines. In A-C, time is shown in min:s. In B and C, the z-plane above the cell where the targeted microbubble was clearly visible was captured before sonication and overlayed on the GCaMP6f images before sonication to indicate the microbubble position. **(D)** Quantification of PI levels as a function of time before and after sonication. **(E)** Comparison of PI levels at 5 min after sonication in non-, reversibly and irreversibly sonoporated cells. Error bars, s.e.m. Arrows point to the reversibly and irreversibly sonoporated cells in B and C, respectively. **(F)** The relative GCaMP6f intensity (F/F_0_) profiles indicate changes in intracellular calcium levels relative to baseline levels in reversibly and non-sonoporated cells before and after sonication. Here, F is the GCaMP6f intensity level at any given time, and F_0_ is the baseline intensity level. Inset: A simple linear regression showing the relationship between intracellular calcium and PI levels in reversibly sonoporated cells. Bands represent the 95% confidence interval. The Pearson coefficient (r) and the p-value are shown. Arrow points to the reversibly sonoporated cell shown in B. In D and F, solid lines and the shaded regions represent the mean and s.e.m., respectively. **(G)** Time profiles of the relative GCaMP6f intensity (F/F_0_) in irreversibly sonoporated cells before and after sonication. The curves end when cells lose their intracellular fluorescence completely, signifying a loss of viability. The time taken for GCaMP6f intensity to reach its minimum value is displayed in the inset. Arrow points to the irreversibly sonoporated cell shown in C. **(H)** Representative images of non- and irreversibly sonoporated cells within a monolayer before and after sonication. **(I-J)** Comparison of the GCaMP6f fluorescence intensity in non- and irreversibly sonoporated cells both before sonication **(I)** and at the time (which varied for each cell as shown in the inset in G) when irreversibly sonoporated cells lost their GCaMP6f fluorescence **(J)**. We selected one non-sonoporated and one reversibly sonoporated cell from each field of view before and after sonication. Error bars, s.e.m. Ultrasound parameters: 267 kHz frequency, 20 μs duration, and ∼110 kPa PNP. Data information: n = 16 reversibly, 6 irreversibly and 16 non-sonoporated cells from N = 3 independent experiments. Statistical analyses were performed using the one-way ANOVA test with Dunnett’s multiple comparisons test in E and the Student’s *t*-test in I and J. Adjusted p-values are shown in E.

In irreversibly sonoporated cells, PI uptake was highest (Fig. 4D, 4E). Akin to reversibly sonoporated cells, intracellular calcium levels rose rapidly and peaked within ∼10 s (Fig. 4G). However, GCaMP6f fluorescence decreased rapidly to significantly lower levels than the baseline value within just 7.9 ± 3.0 min (mean ± s.d.) (Fig. 4G-H). To further characterise this phenomenon, we compared the GCaMP6f fluorescence intensity of non- and irreversibly sonoporated cells in the same frame both before sonication and at the time point when the cells lost their GCaMP6f fluorescence (Fig. 4H-J). The fluorescence intensity was similar in both groups before sonication (Fig. 4I) but was significantly lower in irreversibly sonoporated cells after sonication (Fig. 4J). Moreover, we observed a drastic reduction in the size of reversibly sonoporated cells (Fig. 4C), reminiscent of cell extrusion as reported previously [62,63]. This cascade of events, starting with high PI uptake, followed by the loss of GCaMP6f fluorescence and then cell shrinkage, collectively suggests that these cells underwent irreversible sonoporation and, as a result, lost their viability.

### The mobility of vesicles containing the tight junction protein claudin-5 is reduced to varying degrees in reversibly and irreversibly sonoporated cells

To better understand the morphological changes of the cells undergoing either reversible or irreversible sonoporation, we performed a morphodynamic analysis using an MDCK II cell line that stably expresses the tight junction protein claudin-5 fused to enhanced green fluorescent protein (EGFP-CLDN5 MDCK II) [57]. For this, we located targeted microbubbles in the far-red channel (639 nm) and then imaged EGFP-CLDN5 in the green channel (488 nm) and PI uptake in the red channel (561 nm) in live cells. EGFP-CLDN5 was localised to the cell borders, allowing us to reliably segment the cell boundary (Fig. 5A-B). Consistent with our above observations (Fig. 4), PI uptake after sonication was significantly higher in irreversibly sonoporated cells than in non-or reversibly sonoporated cells (Fig. 5A-C). While the area of both non- and reversibly sonoporated cells remained relatively stable (Fig. 5A-D; Videos S4 and S5), the area of irreversibly sonoporated cells decreased, leading to complete shrinkage (Fig. 5B and 5D; Video S6), confirming the two distinct fates of sonoporated cells observed with our set-up.

**Figure 5.**
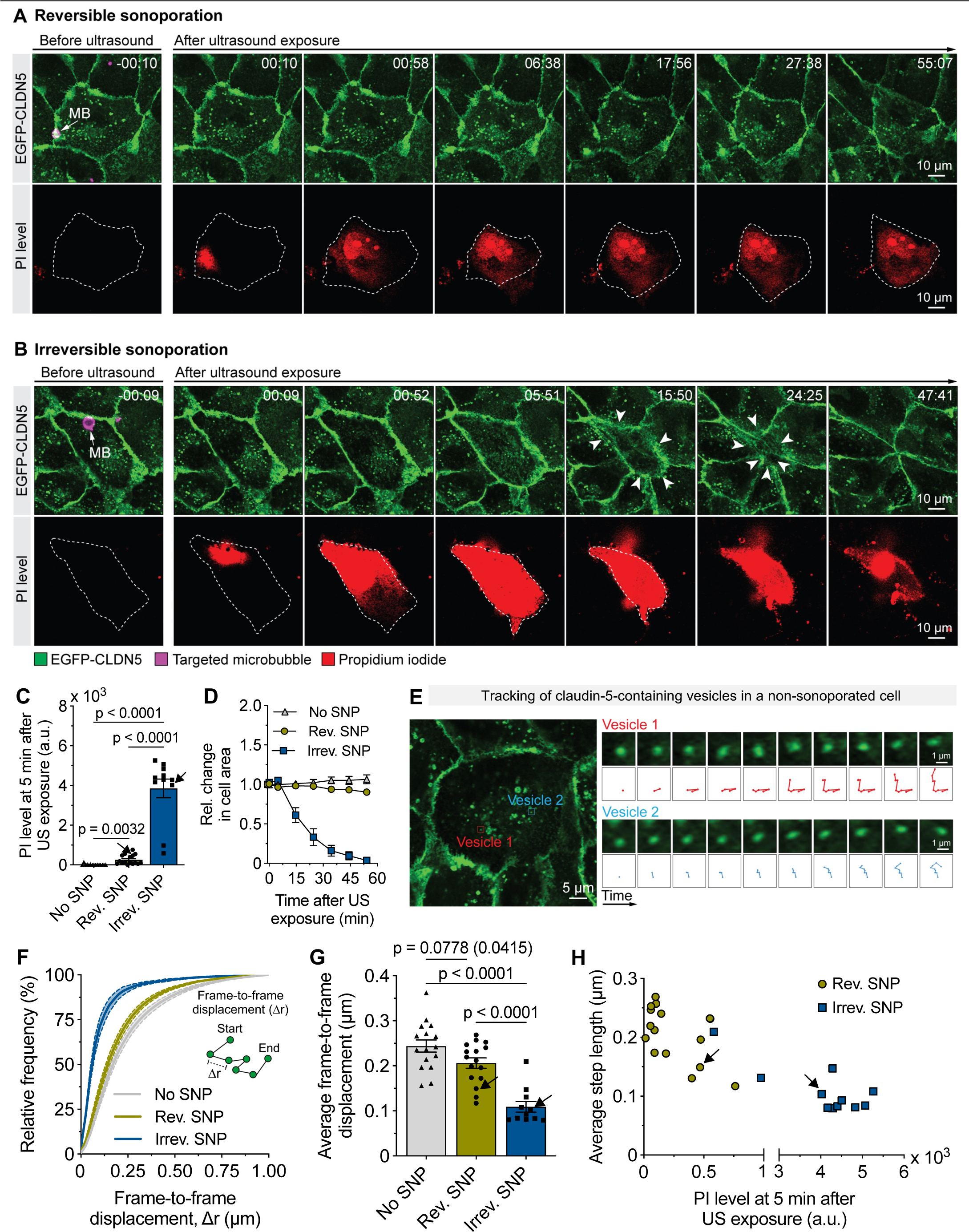
The degree of sonoporation differentially affects CLDN5 vesicle movement dynamics. **(A-B)** Representative time-lapse images of a reversibly **(A)** and irreversibly sonoporated cell **(B)** and corresponding PI levels before and after sonication. EGFP-CLDN5 localises to the cell-cell junctions, demarcating cell boundaries and allowing us to quantify the cell area. Time is shown in min:s. The z-plane above the cell where the targeted microbubble was clearly visible was captured before sonication and overlayed on the EGFP-CLDN5 images before sonication to indicate the microbubble position. **(C)** Comparison of PI levels in reversibly, irreversibly and non-sonoporated cells at 5 min after sonication. Error bars, s.e.m. **(D)** The time profiles of the relative change in the area of reversibly, irreversibly and non-sonoporated cells compared to the area before sonication. **(E)** Snapshot of a non-sonoporated EGFP-CLDN5-MDCK II cell displaying CLDN5-containing vesicles (left). Localisation and tracking of two representative intracellular CLDN5 vesicles over ten consecutive frames at 9.8 s intervals before ultrasound exposure (right). **(F)** The cumulative frequency distribution of the frame-to-frame displacement of CLDN5 vesicles tracked within the first 20 min following sonication. Solid lines and shaded regions represent the mean and s.e.m., respectively. Inset: Schematic of a trajectory and computation of the frame-to-frame displacement, Δr. **(G)** Average displacement of CLDN5 vesicles in reversibly, irreversibly and non-sonoporated cells. Error bars, s.e.m. **(H)** The relationship between the PI levels at 5 min and the average step length of CLDN5 vesicles in reversibly and irreversibly sonoporated cells. Arrows in C, G and H point to the reversibly and irreversibly sonoporated cells shown in A and B. Ultrasound parameters: 267 kHz frequency, 20 μs duration, and ∼110 kPa PNP. Data information: n = 16 non-sonoporated, 16 reversibly and 11 irreversibly sonoporated cells from 4 independent experiments. In C and G, statistical analyses were performed using the one-way ANOVA test with Dunnett’s multiple comparisons test. The adjusted p-values are shown in C and G, with the actual p-values in brackets. In H and I, the bands indicate the 95% confidence interval and the Pearson’s coefficient (r), and the p-values <u>are shown.</u>

In the EGFP-CLDN5-MDCK II cells, we could locate and track the movement of intracellular vesicles containing claudin-5 (CLDN5 vesicles) (Fig. 5E). Serendipitously, our visual inspection suggested that sonoporation appeared to impact the mobility of CLDN5 vesicles (Videos S5 and S6), a process crucial for maintaining cell barrier integrity [64]. We reasoned that quantifying this effect will inform whether sonoporation impacts intracellular vesicular transport. To this end, we calculated the frame-to-frame displacement, Δr, of CLDN5 vesicles. This metric has been widely used to characterise the mobility of intracellular proteins and vesicles [65–67]. We restricted our analysis to the first 20 min following sonication, as thereafter, most irreversibly sonoporated cells shrank drastically. We first computed the cumulative frequency distribution of the frame-to-frame displacement of CLDN5 vesicles. Compared to non-sonoporated cells, in reversibly sonoporated cells, this distribution shifted to the left towards shorter displacement (Fig. 5F). However, this leftward shift was even more pronounced in irreversibly sonoporated cells (Fig. 5F). For statistical comparisons, we estimated the average displacement of CLDN5 vesicles and found that the reduction in the average displacement was greatest in irreversibly sonoporated cells (Fig. 5G-H). These observations indicate that CLDN5 vesicles remain mobile and that the cell size is minimally affected in reversibly sonoporated cells, whereas vesicles stall and the cell size decreases drastically in irreversibly sonoporated cells.

### The microtubule network in reversibly sonoporated cells partially disintegrates and recovers but rapidly collapses in irreversibly sonoporated cells

Given that the intracellular microtubule network is crucial for providing structural support, as well as facilitating vesicle and organelle transport, and maintaining cell morphology, we next asked whether this network is differentially perturbed in reversibly and irreversibly sonoporated cells. To this end, we monitored the microtubule network in EGFP-CLDN5 MDCK II cells using the cell-permeable, fluorogenic, far-red probe SiR-Tubulin, which has high specificity for microtubules. We imaged EGFP-CLDN5 in live cells in the green channel (488 nm), PI uptake in the red channel (561 nm), and SiR-Tubulin signal in the far-red channel (639 nm) (Fig. 6A and 6B).

**Figure 6.**
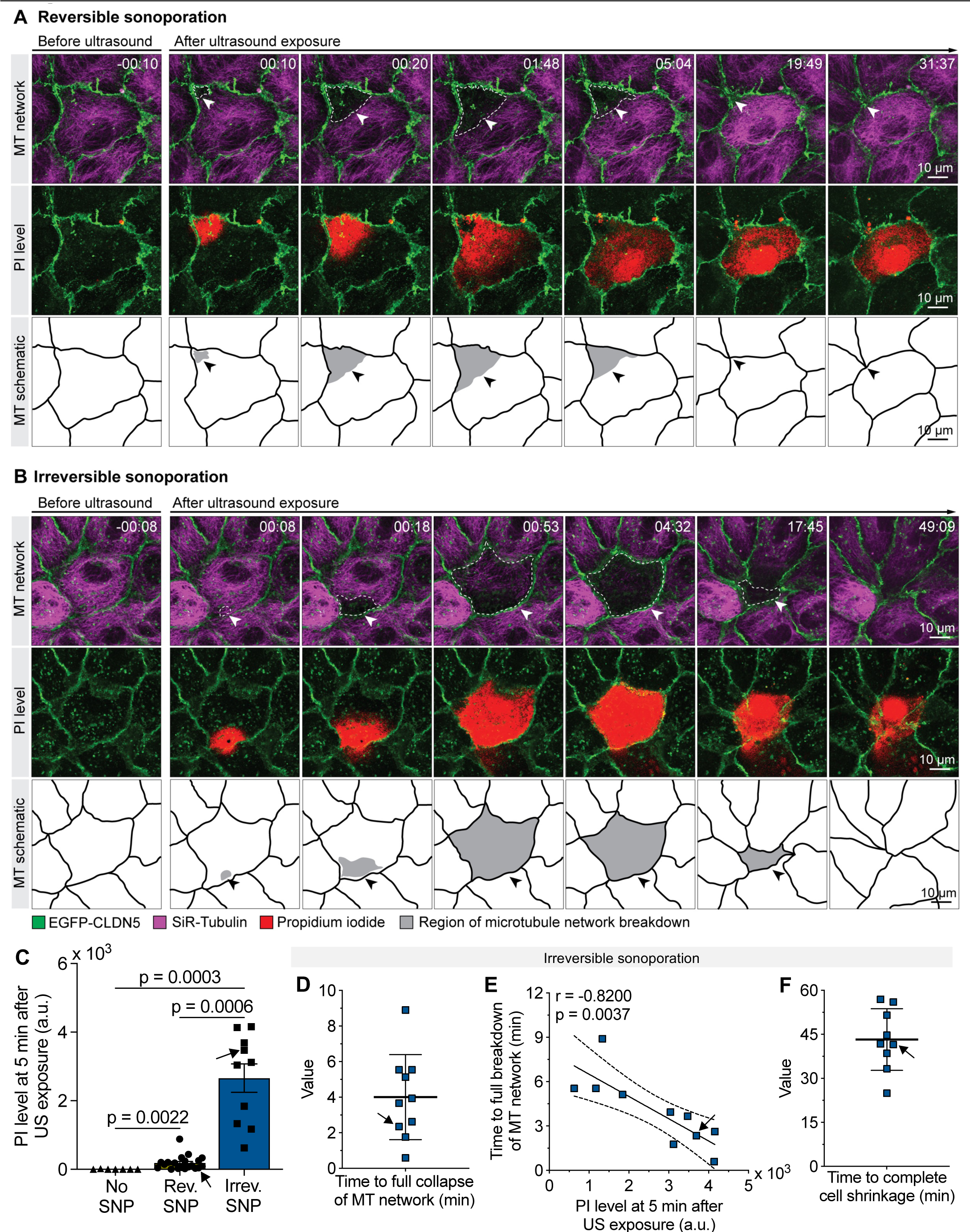
The microtubule network breaks down but recovers in reversibly sonoporated cells, whereas it collapses completely in irreversibly sonoporated cells. **(A-B)** Representative time-lapse images of microtubule breakdown dynamics in a reversibly (A) and irreversibly (B) sonoporated cell with the corresponding PI levels after sonication. EGFP-CLDN5 signals are overlayed onto the SiR-Tubulin and PI signals in the first two rows to demarcate cell boundaries. The schematic highlights the microtubule disrupted area (grey) and the cell boundaries (black line) inferred from the EGFP-CLDN5 and SiR-Tubulin signals. Time is shown in min:s. **(C)** Comparison of the mean PI levels in reversibly, irreversibly and non-sonoporated cells at 5 min after sonication. Error bars, s.e.m. Arrows point to the reversibly and irreversibly sonoporated cells shown in A and B. **(D)** Time taken for the total collapse of the microtubule network in irreversibly sonoporated cells. **(E)** A simple linear regression showing the relationship between PI uptake at 5 min post-sonication and the time until complete microtubule network collapse in irreversibly sonoporated cells. The Pearson’s coefficient (r) and the p-value are shown. **(F)** Time until complete shrinkage of irreversibly sonoporated cells. Arrows in D to F point to the irreversibly sonoporated cell shown in B. Ultrasound parameters: 267 kHz frequency, 20 μs duration, and ∼110 kPa PNP. Data information: n = 12 non-, 20 reversibly and 10 irreversibly sonoporated cells from 4 independent experiments. In C, statistical analyses were performed using the one-way ANOVA test with Dunnett’s multiple comparisons test and the adjusted <u>p-values are shown.</u>

Again, we consistently observed that irreversibly sonoporated cells had the highest levels of PI uptake (Fig. 6C). Crucially, the microtubule network dynamics after ultrasound exposure was strikingly different in reversibly and irreversibly sonoporated cells (Fig. 6A and 6B; Videos S7, S8 and S9). Immediately after sonication, microtubule network breakdown was initiated at the local site of PI uptake in all sonoporated cells. However, whilst the microtubule network recovered in reversibly sonoporated cells (Fig. 6A; Video S8), the entire network collapsed in irreversibly sonoporated cells (Fig. 6B; Video S9), eventually leading to cell shrinkage. In irreversibly sonoporated cells, the network collapse occurred within 4.0 ± 2.4 min (mean ± s.d.) (Fig. 6D-E), well before complete cell shrinkage, which took 43.3 ± 10.5 min (mean ± s.d.) (Fig. 6F). Moreover, the time taken for the entire microtubule network to collapse was inversely correlated with PI uptake (Fig. 6E), suggesting that the degree of sonoporation dictates microtubule network collapse dynamics even within the irreversibly sonoporated cell subpopulation.

In reversibly sonoporated cells, the microtubule breakdown area initially increased and then decreased (Fig. 6A), indicating a partial collapse and subsequent recovery of the network; however, this dynamic behaviour varied significantly across cells exposed to the same ultrasound condition. For example, the maximal microtubule network breakdown area shown for one cell (denoted as cell 1) was 402.8 µm^2^, occurring at ∼1 min, while for another cell (denoted as cell 2), this area was 22.2 µm^2^, occurring at around 30 s (Fig. 7A-B). Not surprisingly, although the microtubule network recovered in both cells, PI uptake levels were markedly different (Fig. 7B inset). We next estimated these metrics for all cells. It took 28.9 ± 18.0 s (mean ± s.d.) to reach the maximal breakdown area (Fig. 7C). The mean maximal microtubule network breakdown area and the mean time for network recovery were 80.4 ± 96.3 µm^2^ and 3.2 ± 2.9 min (mean ± s.d.), respectively (Fig. 7D-E). Additionally, both metrics were positively correlated with PI uptake (Fig. 7F-G), indicating that the degree of sonoporation influenced the dynamics of the microtubule network breakdown and its recovery even within reversibly sonoporated cell subpopulation.

**Figure 7.**
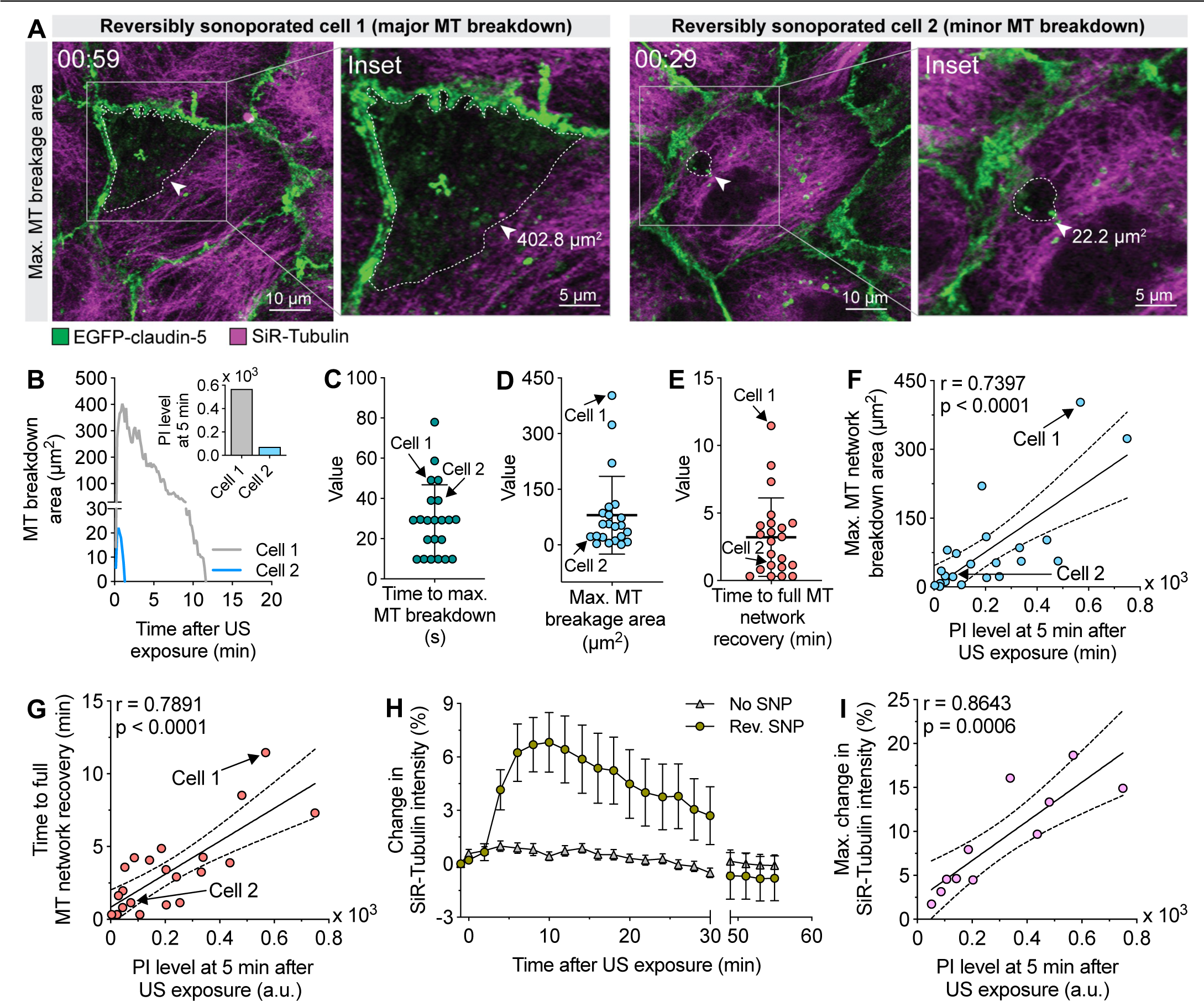
Reversible sonoporation transiently disrupts the microtubule network and triggers a transient increase in tubulin intensity in the intact microtubule network region. **(A)** Examples of microtubule network breakdown in two reversibly sonoporated cells. The degree of breakdown assessed by the maximal area of microtubule breakdown (dashed white line) varies between cells. **(B)** Time profiles of the disrupted microtubule network area in cells 1 and 2. Inset: PI levels 5 min after sonication of cells 1 and 2. (**C)** Time taken for the maximal microtubule network breakdown. (**D)** Maximal microtubule breakdown area. **(E)** Time until microtubule network recovery. In C-E, Error bars, s.d. Arrows point to cells 1 and 2. **(F-G)** Simple linear regression depicting the relationship between the maximal area of microtubule breakdown (F) and the microtubule network recovery time (G) with PI levels in reversibly sonoporated cells. Error bands indicate the 95% confidence intervals. **(H)** Percentage change in tubulin intensity of the unaffected microtubule network region in reversibly sonoporated cells, and of the entire network in non-sonoporation cells. Reversibly sonoporated cells with at least 50 µm^2^ microtubule breakdown area were considered. Error bars, s.e.m. **(I)** Simple linear regression indicating the relationship between the maximum change in tubulin intensity and PI levels in reversibly sonoporated cells. In F, G and I, the Pearson’s coefficient (r) and the p-value are shown. Ultrasound parameters: 267 kHz frequency, 20 μs duration, and ∼110 kPa PNP. Data <u>information: n = 12 non- and 20 reversibly sonoporated cells from 4 independent experiments.</u>

Intriguingly, in reversibly sonoporated cells with extensive microtubule network breakdown, the SiR-tubulin intensity increased in the intact portion of the microtubule network with time (Video S8). We quantified this effect in cells with at least 50 µm^2^ network damage area. We then segmented the cell region with the intact microtubule network and found that the SiR-Tubulin intensity increased in this region, reached a maximum within ∼10 min, and then approached baseline levels within 60 min (Fig. 7H). In contrast, this pattern of SiR-Tubulin intensity changes was not observed in non-sonoporated control cells (Fig. 7H). Furthermore, we found that the maximal increase in SiR-Tubulin intensity positively correlated with PI uptake (Fig. 7I). It is, therefore, conceivable that the free tubulin released in response to the microtubule network disruption diffuses in the cytoplasm and becomes enriched in the microtubule-intact region, leading to increased SiR-Tubulin intensity. These findings highlight the intricate spatiotemporal dynamics of the microtubule network associated with different fates of sonoporated cells.

In summary, sonoporation triggers a cascade of PI uptake, calcium elevation and microtubule network disintegration at a focal site near where the targeted microbubble was attached to the apical cell surface. Then, PI and calcium signals spread throughout the cytoplasm. In reversibly sonoporated cells, the microtubule network recovered rapidly, calcium homeostasis was restored relatively slowly, CLDN5 vesicles retained their mobility, and cell size remained stable. In contrast, in irreversibly sonoporated cells, the entire microtubule network collapsed, CLDN5 vesicles stalled, calcium homeostasis was lost, and cell shrinkage occurred. Our study thus maps the cascade of critical events following sonoporation.

## DISCUSSION

Microbubble-mediated plasma membrane perforation is a dominant mode of action of low-intensity ultrasound treatment that can be exploited for effective drug delivery, provided the molecular and cellular responses in sonoporated cells are fully understood. Here, by developing a custom-built ultrasound device suitable for high-resolution, live-cell imaging of genetically engineered MDCK II cells, we quantified how the spatiotemporal profiles of several bioeffects, ranging from intracellular calcium response to cell morphodynamics, differ in reversibly and irreversibly sonoporated cells (Fig. 8A-B). In both types of sonoporated cells, PI uptake was found to occur locally near a site where microbubbles were targeted via a ligand for the cell surface receptor CD51/61, with the PI signal then spreading throughout the cytoplasm within ∼2 min. This pattern of PI uptake and distribution is consistent with the observations from previous studies using various cell types [29,34,44,55].

**Figure 8.**
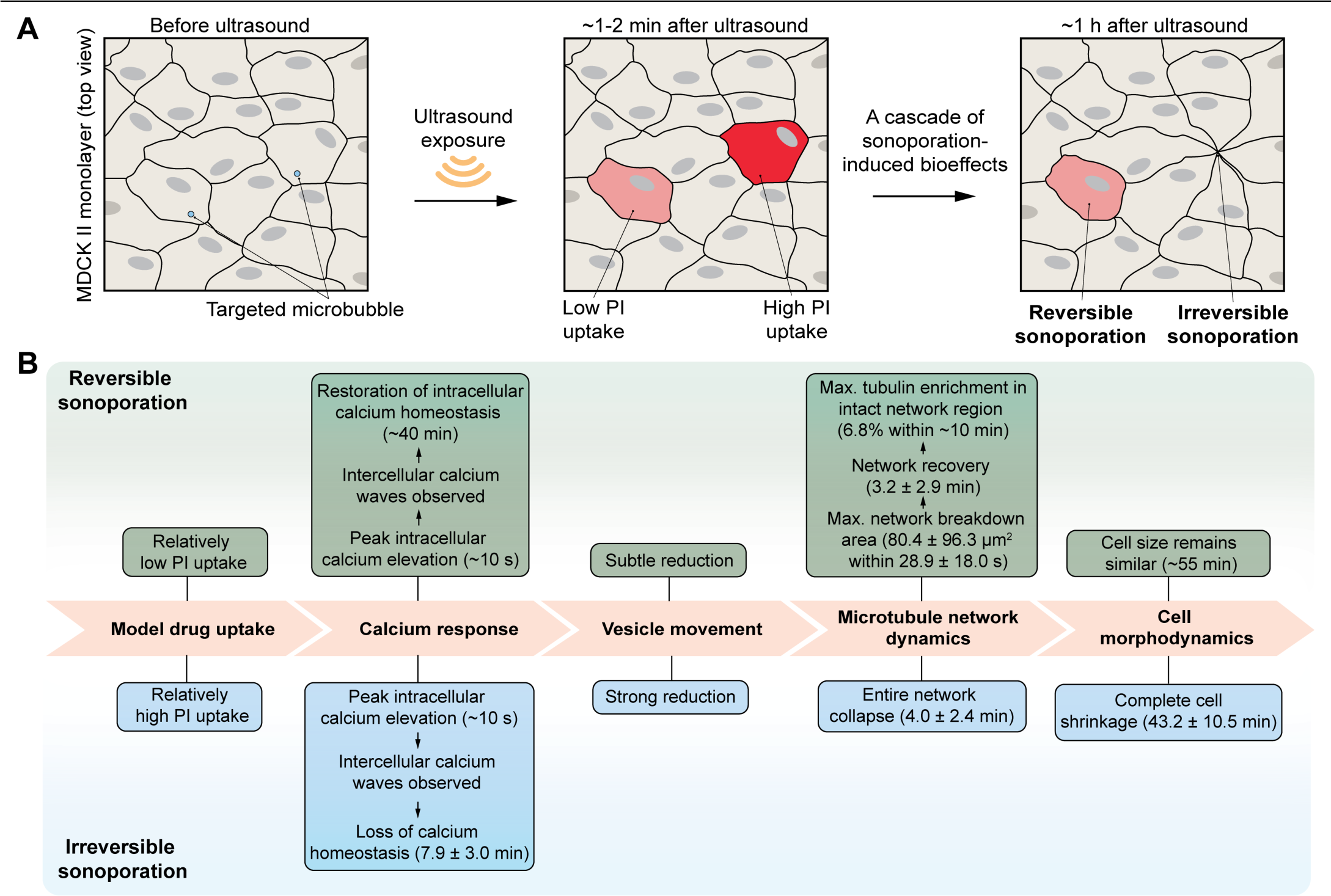
Summary of the cascade of bioeffects differentiating reversibly and irreversibly sonoporated cells. **(A)** A single pulse of ultrasound at a frequency of 267 kHz and a PNP of ∼110 kPa triggers both reversible and irreversible sonoporation within a monolayer in our setup. **(B)** Molecular and cellular responses observed in reversibly and irreversibly sonoporated cells.

In what way do the two types of sonoporated cells differ? In reversibly sonoporated cells, intracellular calcium levels increased, peaked within ∼10 s, and returned to baseline values by ∼40 min. Simultaneously, a microtubule network breakdown occurred, with a maximum breakdown area of 80.4 ± 96.3 µm^2^ (mean ± s.d.) within 28.9 ± 18.0 s (mean ± s.d.). This network recovered within 3.2 ± 2.9 min (mean ± s.d.). The tubulin intensity in the remaining intact microtubule network also increased, peaked by ∼10 min, and then returned to baseline levels. Furthermore, there was a subtle reduction in the movement of vesicles containing the tight junction protein CLDN-5, and the cell size remained relatively similar throughout the entire imaging (55 min). In irreversibly sonoporated cells, intracellular calcium levels peaked within ∼10 s, but the levels reduced considerably below the baseline value within 7.9 ± 3.0 min (mean ± s.d.), indicating a loss of cell viability. Moreover, the entire microtubule network collapsed within 4.0 ± 2.4 min (mean ± s.d.). CLDN-5 vesicle movement was reduced substantially and almost stalled, and the cell volume shrank completely within 43.2 ± 10.5 min (mean ± s.d.). Our study thus offers quantitative insight into the multiscale processes occurring in cells undergoing both reversible and irreversible sonoporation.

Previous imaging-based *in vitro* sonoporation studies have primarily focussed on early bioeffects occurring <15 min following sonoporation [33,34,36,39,42–45,55]. Our setup allowed us to maintain the cells in culture medium at 37° C with appropriate levels of CO_2_ and humidity for extended periods, up to an hour. Using cells without microbubbles attached (non-sonoporated cells) as an internal control, we minimised the impact of any potential artefacts due to imaging conditions. This approach not only enabled us to image the biological responses up to 1 h post-sonification but also to relate the biological changes to reversible or irreversible sonoporation, thereby providing a platform for identifying critical bioeffects that determine the fates of sonoporated cells. We first observed that all sonoporation events, defined by the uptake of the model drug PI, were accompanied by fluctuations in intracellular calcium levels, consistent with previous reports [32–38]. However, we found that the long-term dynamics of the calcium response were markedly different in reversibly and irreversibly sonoporated cells. Moreover, in agreement with previous studies [32–34,36,37,68], we found that intercellular calcium waves spread to the cells surrounding the sonoporated cells. Intriguingly, calcium levels remained elevated in the reversibly sonoporated cells for ∼40 min, indicating a slow restoration of normal homeostatic concentrations. Future work is needed to identify the molecular players involved in the intra- and intercellular calcium responses and to determine whether other cellular functions of reversibly sonoporated cells, such as cell proliferation, are affected over the timescale of several hours to days.

The microtubule network is integral to any cell’s cytoskeleton, facilitating vesicular transport and providing structural integrity [69,70]. To our knowledge, two studies in HeLa cells have examined the short-term impact of sonoporation on the microtubule network [44,71]. Their observations were limited to up to 3 min post-sonication, reporting a reduction in fluorescence intensity of GFP-labelled α-tubulin [44,71]. However, the long-term consequences of sonoporation on microtubule network recovery and integrity, as well as cell fate, remained unknown. Our setup allowed us to resolve the spatial and temporal dynamics of the microtubule network for up to 1 h post-sonoporation. Specifically, we found that an initial slow disassembly and partial breakdown of the microtubule network, followed by full network recovery, led to reversible sonoporation, whereas a fast disassembly and full collapse of the microtubule network led to cell shrinkage and irreversible sonoporation. Notably, the reduction in CLDN5 vesicle movement was more pronounced in the irreversibly sonoporated cells than in the reversibly sonoporated cells, suggesting that vesicular movement and microtubule integrity are interlinked in sonoporated cells. An exciting future direction would be to assess the association between these changes and transcellular drug transport.

Optical imaging studies examining sonoporation have, thus far, predominantly used MHz frequencies, which is in the range typically used for ultrasound imaging [32–38,41,42,44,49,55]. However, for brain-targeted treatment strategies in humans, as skull attenuation positively correlates with ultrasound frequency, kHz frequencies are being used [50,52,72]. Here, we provide supporting evidence that a single-shot ultrasound pulse at 267 kHz and a pressure of ∼110 kPa triggers both reversible and irreversible sonoporation in a monolayer of MDCK II cells. An important focus for future research will be to examine whether different sonication parameters, such as acoustic pressure, frequency and pulse duration, elicit distinct spatiotemporal profiles of downstream bioeffects.

Prior *in vitro* studies using primary human umbilical vein endothelial cells have reported the opening of cell-cell contacts following sonoporation [29,43,48,55]. In our experiments, we observed overt changes in the cell-cell contact regions of irreversibly sonoporated cells. In contrast, in reversibly sonoporated cells, we did not detect any cell-cell contact opening. Given that the integrity of cell-cell contacts before treatment is a critical determinant of sonoporation-induced contact opening [43,55], it is possible that under our experimental conditions, the confluent MDCK II cells maintained high levels of contact integrity, which prevented contact openings in reversibly sonoporated cells. Another possibility is that perhaps we may not have sampled cells that were sonoporated sufficiently enough to cause contact opening without inducing irreversible sonoporation. It is also conceivable that sonoporation-induced contact opening response is cell-type-dependent. Further work, particularly *in vivo*, is needed with different cell types to better understand the links between cell-cell contact opening and the different long-term fates of sonoporated cells.

We recognise the limitations of our study. First, our analysis was restricted to MDCK II epithelial cells. Although we believe that downstream bioeffects, such as microtubule network breakdown, recovery and collapse dynamics, are likely shared across cell types, we note that the degree of sonoporation varies across cell types [73], and therefore, the kinetics of bioeffects may similarly differ across cell types. Since microbubbles interact with different cell types, such as endothelial and cancer cells, depending on the therapeutic context, future studies could include these cell types to ascertain the broader relevance of our findings. Moreover, employing 3D tumour spheroid or 3D microvascular blood-brain barrier models in future studies may overcome the limitations of 2D monolayer cell cultures used in our study. Second, although PI has been widely used in sonoporation studies [29,31,32,36–38,41–44,48,49,74,75], it can be toxic to cells over time. Therefore, future long-term, live-cell imaging studies could consider alternative fluorescently labelled molecules that are less toxic. Third, because of the standing waves generated by acoustic reflection at the bottom of the dish, the actual pressure experienced by the cells likely differs from what is measured in the free field. Additionally, further investigation is needed to assess the extent to which standing waves impact microbubble dynamics and subsequent sonoporation events. Finally, our imaging duration was up to ∼1 h post-sonoporation, sufficient to classify reversible and irreversible sonoporation based on cell morphological changes. A recent study showed that the long-term proliferation and restoration capabilities of reversibly sonoporated cells are altered over several hours to days [38]. Future work is needed to investigate how other functions relevant to drug delivery, such as transcytosis, are altered long-term in reversibly sonoporated cells. Another key area for future research is exploring the link between ultrasound-driven microbubble dynamics [8,76], which our study did not quantify, and sonoporation events.

In summary, we quantified the differences in vesicle movement, intracellular calcium response, microtubule network dynamics, PI uptake, and cell morphodynamics that distinguish reversible from irreversible sonoporation in an *in vitro* model. Our study not only improves our knowledge of sonoporation-induced bioeffects occurring at different spatial and temporal scales but also provides a platform to identify targetable bioeffects to manipulate the fate of sonoporated cells.

## METHODS

### Cell culture

MDCK II cells (ECACC #00062107), including stable cell lines expressing GCaMP6f and EGFP-CLDN5, were maintained in Minimum Essential Medium (MEM) (Sigma; Cat# M4655) supplemented with 5% foetal bovine serum (FBS, Bovogen, Australia; Cat# SFBS-FR), 100 U/ml penicillin, and 100 µg/ml streptomycin (Gibco; Cat# 15140122). Cells were grown at 5% CO_2_ in 37°C humidified air. For imaging, cells were cultured in Fluorobrite^TM^ Dulbecco’s Modified Eagle Medium (FB-DMEM, Gibco; Cat# A1896701) supplemented with 5% FBS, 1% GlutaMax (Gibco; Cat# 35050061), 100 U/ml penicillin, and 100 µg/ml streptomycin. For all imaging experiments, cells were seeded at a density of 1.4 x 10^6^ cells/dish in 35 mm glass-bottom imaging dishes (Ibidi, Gräfelfing, Germany; Cat# 81158) and allowed to grow to confluence over 48 h.

### Cell line generation

#### Construction of the GCaMP6f lentiviral vector

To generate the lentiviral vector Ef1α-GCaMP6f, the GCaMP6f coding sequence was amplified by polymerase-chain reaction (PCR) from pGP-CMV-GCaMP6f (Addgene #40755) using primers encoding the *BamHI* and *MluI* restriction enzyme sites (forward primer: AATTAAGGATCCGC-CACCATGGGTTCTCATC; reverse primer: AATTAAACGCGTGCGGCC-GCTCACTTCGCTGTC). The GCaMP6f fragment was then digested using the restriction enzymes BamHI-HF and MluI-HF (NEB) and ligated into the empty lentiviral vector pLV-EF1α using T4 DNA ligase (NEB). Plasmids were purified using the Nucleobond Xtra EF Midi Kit (Macherey-Nagel) and confirmed by Sanger sequencing.

#### Transfection and lentivirus production

Lenti-X 293T cells (Takara Bio) were seeded in six-well plates and transfected 24 h later at 70% confluency. Before transfection, the medium was replaced with DMEM supplemented with 10% FBS and 10 mM HEPES (Thermo Fisher), and cells were transfected with TransIT-VirusGEN (Mirus) according to the manufacturer’s instructions using 0.23 µg pMD2.G (Addgene #12259), 0.77 µg PsPAX2 (Addgene #12260), and 1 µg Ef1α-GCaMP6f per well. Virus-containing medium was collected at 72 h post-transfection, passed through a 0.45 µm filter and mixed with an in-house lentiviral concentrator solution (4X solution of 40% PEG6000, 1.2 M NaCl and 1X phosphate-buffered saline (PBS)). Viral supernatants were pooled and precipitated at 4°C for >1 h, spun at 4°C for 45 min at 1,500 *g* (Microfuge 22R, Beckman Coulter), and concentrated 100-fold by resuspending the viral pellets in Opti-MEM medium.

#### Infection of MDCKII cells and establishment of the monoclonal line

MDCK II cells were seeded in six-well plates and infected with 15 µl of concentrated lentivirus when cells reached approximately 50% confluence. Individual cells were sorted approximately one week post-infection using a BD FACSAria Cell Sorter into 96-well plates based on a gating strategy to select the brightest 10% of GCaMP6f-expressing cells. The monoclonal line with the brightest and most uniform expression of GCaMP6f was chosen for further use.

### Microbubble preparation

A mixture of lipids comprising of 1,2-distearoyl-sn-glycero-3-phosphocholine (DSPC; 90 mol%; Avanti; Cat# 850365P), 1,2-distearoyl-sn-glycero-3-phosphoethanolamine-N-[carboxy(polyethylene glycol)2000] (DSPE-PEG(2000), 8.4 mol%, Avanti; Cat# 880128P) and 1,2-distearoyl-sn-glycero-3-phosphoethanolamine-N-[biotinyl(polyethylene glycol)-2000] (DSPE-PEG(2000)-biotin, 1.6 mol%, BroadPharm; Cat# BP-22723) was dissolved in 1.5 ml of chloroform (Sigma) in a glass beaker. The chloroform solvent was evaporated under vacuum (Mivac Quattro, Genevac Ltd, Ipswich, Suffolk, United Kingdom) at 22-28°C for 30 min and the dried lipid deposit was rehydrated with a mixture of 10% glycerol (Millipore; Cat# 356350) and 10% propylene glycol (Sigma; Cat# P4347) in 1X PBS to achieve a concentration of 1 mg lipid/ml. The rehydrated lipid mixture was then heated to 55°C in a sonicating water bath until the lipid deposit was fully dissolved. To fluorescently label the lipid coating, the lipid dye DiD (1,1′-dioctadecyl-3,3,3′,3′-tetramethylindodicarbocyanine perchlorate, 1 mol%; Invitrogen) was added to the lipid mixture during the rehydration step before it was heated and sonicated. The solution was cooled, then aliquoted into 1.5 ml glass scintillation vials and the air headspace was replaced with octafluoropropane (C_3_F_8_) (Arceole) using a 27G needle. Microbubbles were generated by agitation of the vial at 4,000 rpm for 45 s using a dental amalgamator. Before use, microbubbles were washed three times with filtered 1X PBS in a 5 ml Luer-lock syringe through centrifugation at 400 *g* for 1 min using a bucket-rotor centrifuge (Allegra X30-R, SX4400 rotor, Beckman Coulter) to remove excess lipids. Non-targeted microbubbles were obtained after these three wash steps. Microbubble concentration was determined after the final washing step with a Coulter Counter Multisizer 4e (30 µm aperture; Beckman Coulter) and was at least 10^9^ microbubbles/ml.

To target microbubbles to the cell surface of MDCK II cells, they were functionalised with an antibody against CD51/61 using the biotin-streptavidin bridging method. This method involves linking a biotinylated antibody to the biotinylated lipid microbubble shell through streptavidin. Firstly, 6 x 10^8^ microbubbles were incubated on ice for 30 min with 60 µg of streptavidin (1 mg/ml stock concentration in ultrapure water, Invitrogen; Cat# 434301) and washed once by centrifugation at 400 *g* for 1 minute in a 5 ml Luer-lock syringe filled with 1x PBS. Streptavidin-conjugated microbubbles were counted after washing to ensure microbubble viability. Then, streptavidin-conjugated microbubbles were incubated on ice for 30 min with 6 µg of biotinylated anti-CD51/61 antibody (BioLegend; Cat# 304412) before a final wash step, and a final count was performed to obtain the final targeted microbubble concentration and size distribution. When SiR-Tubulin dye was added to the culture, targeted microbubbles without DiD labelling were generated and used.

### Evaluation of PI uptake

PI, a fluorogenic, membrane-impermeable agent, was used as a model drug and an indicator of sonoporation [77]. Upon sonoporation, PI diffuses into the cell through the membrane pore and binds to RNA and DNA in the cytoplasm and nucleus [77], where it forms complexes that fluoresce with an emission maximum of 617 nm and excitation maximum of 535 nm. PI has been used in numerous studies to evaluate membrane perforation due to sonoporation [32,38,55]. PI (Invitrogen; Cat# P3566) was added at a final concentration of 100 µM to the culture dish and mixed well before sonication. To ensure homogeneous mixing, PI was first added to 1 ml of medium in a 1.5 ml Eppendorf tube, mixed well by pipetting, and then added to a further 1 ml of medium in the culture dish to a final volume of 2 ml.

### Microtubule staining

Confluent MDCK II cells were incubated with culture medium containing 500 nM SiR-Tubulin (Spirochrome, Switzerland) supplemented with 10 µM verapamil for 2-6 h before imaging. Immediately before loading of targeted microbubbles, SiR-Tubulin-containing media was removed. Following the attachment of microbubbles, cells were washed twice with warm culture medium, and fresh FB-DMEM without SiR-Tubulin was added. For each independent experiment, we prepared three to four cell culture dishes and treated them with SiR-Tubulin and verapamil for at least 2 h until imaged. As each dish required approximately one hour for imaging, treatment durations varied between 2 and 6 h across the dishes.

### Free-field characterisation of ultrasound transducer

The acoustic profile and focus of a single-element, circular 267 kHz transducer (Fraunhofer IBTM, Germany) of 18 mm diameter were characterised using a transducer scanning apparatus in a water tank (Acoustic Intensity Measurement System, Onda Corp., U.S.A.) filled with degassed water (Fig. 1). The transducer was driven by a TPO with an impedance of 50 0. As the electrical impedance of the transducer is 399 0, we added an RF impedance transformer to minimise the impedance mismatch. The transducer was mounted to a motorised xyz translator coupled to a 1 mm hydrophone (HNR-1000, Onda Corp., U.S.A.), and the transducer surface was submerged in degassed water. Ultrasound waveforms received by the hydrophone were recorded by a computer-controlled oscilloscope (PicoScope, 5244A) in synchronisation with ultrasound excitation. While the hydrophone was scanning in the x-y-z directions at step sizes of 0.6 mm in the z-axis and 0.45 mm in the x- and y-axes, ultrasound pulses were generated by the transducer driven by a TPO system (TPO^TM^, Sonic Concepts Inc., U.S.A.) via a radiofrequency (RF) impedance transformer (T1K-7A, T&C Power Conversion Inc., U.S.A.). The scan covered a z-depth range from 2 to 30 mm away from the transducer surface.

### Ultrasound exposure

The ultrasound device was positioned around the Zeiss incubation insert of a laser-scanning confocal microscope (LSM 980 NLO Airyscan 2, Carl Zeiss GmbH, Germany), which contained the culture dish. Once the device was in position, a customised CO_2_ cover with an opening for the transducer and coupling cone was placed over the incubation insert. Then, using the gross height translator on the ultrasound device, the transducer, immersed in a water-filled coupling cone, was lowered through the CO_2_ cover such that the coupling cone membrane interfaced with the medium within the dish. The height was adjusted such that the distance from the transducer surface and the bottom of the 35 mm imaging dish, where the cells were cultured, was 20 mm. With the CO_2_ cover and incubation insert, cells were maintained at 37°C and 5% CO_2_ throughout the imaging duration. To expose the cells to ultrasound, the 267 kHz single-element transducer was driven by the TPO system and RF impedance transformer to produce a 20 µs single-pulse sinusoidal signal at an average PNP of ∼110 kPa (peak power input = 4 W). We note that standing waves would be generated due to the reflection of sound waves at the bottom of the dish containing cells and that the actual pressure experienced by the cells will be different from the pressure levels measured in the free field (Fig. 1). However, this uncertainty in the estimates of actual pressure will not impact our findings, as we are primarily interested in understanding and differentiating the downstream bioeffects in reversibly and irreversibly sonoporated cells.

### Live-cell imaging

Cells were visually inspected to ascertain full confluence prior to experiments. Before imaging, the culture medium was completely aspirated from the dish, and 20 µl of medium containing targeted microbubbles was added directly on top of the cells. The dish was inverted and placed in an incubator (37°C, 5% CO_2_) for 10 min. Twenty microlitres of medium were sufficient to form a thin layer of liquid over the cells such that the targeted microbubbles can float and bind to the cells, while not dripping when the dish was inverted. The dish was reverted after 10 min and washed once with a warm culture medium to remove unbound or excess microbubbles before adding the medium containing PI.

All imaging was performed using a Zeiss C-Apochromat 40x/1.2 W Korr FCS water-immersion objective on a confocal laser-scanning microscope (LSM 980 NLO Airyscan 2, Carl Zeiss GmbH, Germany) built around an Axio Observer 7 body, equipped with an Airyscan 2 super-resolution detector, and controlled by Zen Blue Software. The working distance of the objective was 280 µm, with a field of view of 211.5 x 211.5 µm (4,084 x 4,084 pixels).

To highlight the bioeffects occurring in sonoporated cells, this field of view was cropped to 51.79 x 51.79 µm (1,000 x 1,000 pixels) for GCaMP6f-expressing cells and 67.33 x 67.33 µm (1,300 x 1,300 pixels) for EGFP-CLDN5-expressing cells, such that the sonoporated cell of interest and several neighbouring cells were visible. Imaging was performed in the green, red, and far-red channels using the Multiplex mode with the SR-8Y acquisition strategy, and images were deconvolved using automatic Airyscan processing prior to image analysis. The improved illumination and detection readout using these modes allow for reduced illumination intensities at the specimen plane, resulting in reduced phototoxicity [78,79]. GCaMP6f and EGFP-CLDN5 were both excited at 488 nm (green, detected at 495-550 nm) with laser powers of 0.07 and 0.08 kW/cm^2^, respectively, and detector gain of 750 V. PI was excited at 561 nm (red, detected at 573-627 nm) with a laser power of 0.04 kW/cm^2^ and detector gain of 850 V. DiD and SiR-Tubulin were both excited at 639 nm (far-red, detected at 655-735 nm) with laser powers of 0.01 kW/cm^2^ and 0.17-0.34 kW/cm^2^ respectively, and detector gains of 750 V and 850-875 V, respectively. Each field of view was scanned bidirectionally, and each channel was excited sequentially such that there was no spectral overlap between them. Definite Focus was implemented at each time point to ensure cells remained in focus. Each frame was acquired every ∼8 to 9.8 s to capture.

Cells were imaged for at least 1 min before sonication to record the baseline values. The end of the frame before delivering a single 20 µs pulse of ultrasound was defined as time *t* = 0. The cellular response following microbubble insonification was recorded for at least 55 min after ultrasound exposure. One field of view, comprising several single microbubbles attached to different cells, was imaged per dish. The field of view was chosen based on the following criteria: i) the region of the monolayer within the field of view was fully confluent, and ii) the presence of several cells with a single targeted microbubble attached to them. Cells in the field of view without microbubbles attached to them did not undergo sonoporation and thus served as an internal control.

### Vesicle tracking

CLDN5-containing vesicle tracking was performed with the TrackMate plugin in ImageJ [80]. Videos were first cropped such that the region of interest consisted of only the intracellular CLDN5 vesicles. The LoG detector with an estimated object diameter of 1.5 µm and quality threshold of 15 was applied along with median filter pre-processing, and sub-pixel localisation was used. Following this, the simple LAP tracker with a maximum particle linking distance and gap-closing maximum distance of 1 µm was used. The maximum gap size was set to 2 frames. All parameters were adjusted empirically to accurately localise individual vesicles within the cell. We first calculated the frame-to-frame displacement and then computed the empirical cumulative frequency distribution of the displacements of vesicles using the ECDF tool in MATLAB (MathWorks, Natick, Massachusetts, U.S.A.) [81,82].

### Data analysis

Image analysis was performed by selecting individual cells of interest from the entire field of view (211.5 x 211.5 µm). ROIs were drawn around cells of interest at various time points to measure cell area, PI uptake, and response to sonoporation. In all our experiments, cells with PI uptake and a stable size for 55 min post-sonication were classified as reversibly sonoporated cells, while those with PI uptake and a significant size reduction within 55 min were deemed irreversibly sonoporated cells. For reversibly sonoporated cells, the regions of interest could be drawn for the entire imaging duration (∼1 h). However, for irreversibly sonoporated cells, regions of interest were only drawn up until the cell volume shrank completely. For GCaMP6f-expressing cells, cell outlines were manually drawn based on the natural variation of the GCaMP6f signal across cells and the PI intensity within sonoporated cells. For EGFP-CLDN5-expressing cells, the green fluorescence signal at the cell-cell boundaries was sufficiently clear to outline the cell manually. PI uptake levels were computed by subtracting the initial PI intensity measured in the first frame from the PI intensity measured in all subsequent frames. All image analyses were performed using ImageJ version 1.53c (ImageJ Software, Bethesda, Maryland, U.S.A.). Images were acquired with a Zeiss Airyscan 2 detector module in 16-bit mode, producing an intensity level range of 0-65,535. For visualisation purposes, images in the figure are displayed using a fixed intensity range. For instance, the intensity ranges used for GCaMP6f and PI images in Fig. 4 are 9,700-18,000 and 10,200-11,600, respectively. All statistical analyses were carried out with GraphPad Prism version 10.4.1 (GraphPad Software, Boston, Massachusetts, U.S.A.). Bleach correction was performed on GCaMP6f and SiR-Tubulin signal using Huygens Professional deconvolution software before intensity analyses.

## ACKNOWLEDGEMENTS

The imaging was performed at the Queensland Brain Institute Advanced Microscopy Facility, supported by the Australian Government through the Australian Research Council LIEF grant (LE130100078). We thank Dr Adam Briner and Dr Liyu Chen for their assistance with the generation of the plasmid constructs and stable cell lines. We acknowledge the support by the Yulgilbar Foundation to J.L. and the National Health and Medical Research Council of Australia to J.G. (GNT1176326) and P.P. (GNT2026929).

## COMPETING INTERESTS

The authors declare that no conflicts of interest exist.

## Notes

### Competing Interest Statement

The authors have declared no competing interest.

## REFERENCES

[1] A. Bouakaz, J. Michel Escoffre, From concept to early clinical trials: 30 years of microbubble-based ultrasound-mediated drug delivery research, Adv. Drug Deliv. Rev. 206 (2024) 115199. 10.1016/j.addr.2024.115199.

[2] S.M. Chowdhury, L. Abou-Elkacem, T. Lee, J. Dahl, A.M. Lutz, Ultrasound and microbubble mediated therapeutic delivery: Underlying mechanisms and future outlook, J. Controlled Release 326 (2020) 75–90. 10.1016/j.jconrel.2020.06.008.

[3] K. Kooiman, S. Roovers, S.A.G. Langeveld, R.T. Kleven, H. Dewitte, M.A. O’Reilly, J.-M. Escoffre, A. Bouakaz, M.D. Verweij, K. Hynynen, I. Lentacker, E. Stride, C.K. Holland, Ultrasound-Responsive cavitation nuclei for therapy and drug delivery, Ultrasound Med. Biol. 46 (2020) 1296–1325. 10.1016/j.ultrasmedbio.2020.01.002.

[4] A. Jangjou, A.H. Meisami, K. Jamali, M.H. Niakan, M. Abbasi, M. Shafiee, M. Salehi, A. Hosseinzadeh, A.M. Amani, A. Vaez, The promising shadow of microbubble over medical sciences: from fighting wide scope of prevalence disease to cancer eradication, J. Biomed. Sci. 28 (2021) 49. 10.1186/s12929-021-00744-4.

[5] A. Rix, A. Curaj, E. Liehn, F. Kiessling, Ultrasound microbubbles for diagnosis and treatment of cardiovascular diseases, Semin. Thromb. Hemost. 46 (2020) 545–552. 10.1055/s-0039-1688492.

[6] J.H. Song, A. Moldovan, P. Prentice, Non-linear acoustic emissions from therapeutically driven contrast agent microbubbles, Ultrasound Med. Biol. 45 (2019) 2188–2204. 10.1016/j.ultrasmedbio.2019.04.005.

[7] I. Lentacker, I. De Cock, R. Deckers, S.C. De Smedt, C.T.W. Moonen, Understanding ultrasound induced sonoporation: definitions and underlying mechanisms, Adv. Drug Deliv. Rev. 72 (2014) 49–64. 10.1016/j.addr.2013.11.008.

[8] S. Roovers, T. Segers, G. Lajoinie, J. Deprez, M. Versluis, S.C. De Smedt, I. Lentacker, The role of ultrasound-driven microbubble dynamics in drug delivery: from microbubble fundamentals to clinical translation, Langmuir ACS J. Surf. Colloids 35 (2019) 10173–10191. 10.1021/acs.langmuir.8b03779.

[9] S. Mehier-Humbert, T. Bettinger, F. Yan, R.H. Guy, Plasma membrane poration induced by ultrasound exposure: implication for drug delivery, J. Control. Release Off. J. Control. Release Soc. 104 (2005) 213–222. 10.1016/j.jconrel.2005.01.007.

[10] Y. Hu, J.M.F. Wan, A.C.H. Yu, Membrane perforation and recovery dynamics in microbubble-mediated sonoporation, Ultrasound Med. Biol. 39 (2013) 2393–2405. 10.1016/j.ultrasmedbio.2013.08.003.

[11] J. Rich, Z. Tian, T.J. Huang, Sonoporation: past, present, and future, Adv. Mater. Technol. 7 (2022) 2100885. 10.1002/admt.202100885.

[12] Y. Yang, Q. Li, X. Guo, J. Tu, D. Zhang, Mechanisms underlying sonoporation: interaction between microbubbles and cells, Ultrason. Sonochem. 67 (2020) 105096. 10.1016/j.ultsonch.2020.105096.

[13] B.D.M. Meijering, L.J.M. Juffermans, A. van Wamel, R.H. Henning, I.S. Zuhorn, M. Emmer, A.M.G. Versteilen, W.J. Paulus, W.H. van Gilst, K. Kooiman, N. de Jong, R.J.P. Musters, L.E. Deelman, O. Kamp, Ultrasound and microbubble-targeted delivery of macromolecules is regulated by induction of endocytosis and pore formation, Circ. Res. 104 (2009) 679–687. 10.1161/CIRCRESAHA.108.183806.

[14] I. De Cock, E. Zagato, K. Braeckmans, Y. Luan, N. de Jong, S.C. De Smedt, I. Lentacker, Ultrasound and microbubble mediated drug delivery: acoustic pressure as determinant for uptake via membrane pores or endocytosis, J. Control. Release Off. J. Control. Release Soc. 197 (2015) 20–28. 10.1016/j.jconrel.2014.10.031.

[15] J. Wang, Z. Li, M. Pan, M. Fiaz, Y. Hao, Y. Yan, L. Sun, F. Yan, Ultrasound-mediated blood-brain barrier opening: An effective drug delivery system for theranostics of brain diseases, Adv. Drug Deliv. Rev. 190 (2022) 114539. 10.1016/j.addr.2022.114539.

[16] K.-H. Song, B.K. Harvey, M.A. Borden, State-of-the-art of microbubble-assisted blood-brain barrier disruption, Theranostics 8 (2018) 4393–4408. 10.7150/thno.26869.

[17] D. Przystupski, M. Ussowicz, Landscape of Cellular Bioeffects Triggered by Ultrasound-Induced Sonoporation, Int. J. Mol. Sci. 23 (2022) 11222. 10.3390/ijms231911222.

[18] J. Tu, A.C.H. Yu, Ultrasound-mediated drug delivery: sonoporation mechanisms, biophysics, and critical factors, BME Front. 2022 (2022) 9807347. 10.34133/2022/9807347.

[19] J. Deprez, G. Lajoinie, Y. Engelen, S.C. De Smedt, I. Lentacker, Opening doors with ultrasound and microbubbles: Beating biological barriers to promote drug delivery, Adv. Drug Deliv. Rev. 172 (2021) 9–36. 10.1016/j.addr.2021.02.015.

[20] M. Todorova, V. Agache, O. Mortazavi, B. Chen, R. Karshafian, K. Hynynen, S. Man, R.S. Kerbel, D.E. Goertz, Antitumor effects of combining metronomic chemotherapy with the antivascular action of ultrasound stimulated microbubbles, Int. J. Cancer 132 (2013) 2956–2966. 10.1002/ijc.27977.

[21] A.K.W. Wood, R.M. Bunte, H.E. Price, M.S. Deitz, J.H. Tsai, W.M.-F. Lee, C.M. Sehgal, The disruption of murine tumor neovasculature by low-intensity ultrasound-comparison between 1- and 3-MHz sonication frequencies, Acad. Radiol. 15 (2008) 1133–1141. 10.1016/j.acra.2008.04.012.

[22] Y. Engelen, D.V. Krysko, I. Effimova, K. Breckpot, M. Versluis, S. De Smedt, G. Lajoinie, I. Lentacker, Optimizing high-intensity focused ultrasound-induced immunogenic cell-death using passive cavitation mapping as a monitoring tool, J. Control. Release Off. J. Control. Release Soc. 375 (2024) 389–403. 10.1016/j.jconrel.2024.09.016.r

[23] G. Leinenga, J. Götz, Scanning ultrasound removes amyloid-β and restores memory in an Alzheimer’s disease mouse model, Sci. Transl. Med. 7 (2015) 278ra33. 10.1126/scitranslmed.aaa2512.

[24] G. Leinenga, J. Götz, Safety and efficacy of scanning ultrasound treatment of aged APP23 mice, Front. Neurosci. 12 (2018) 55. 10.3389/fnins.2018.00055.

[25] A.R. Rezai, M. Ranjan, M.W. Haut, J. Carpenter, P.-F. D’Haese, R.I. Mehta, U. Najib, P. Wang, D.O. Claassen, J.L. Chazen, V. Krishna, G. Deib, Z. Zibly, S.L. Hodder, K.C. Wilhelmsen, V. Finomore, P.E. Konrad, M. Kaplitt, Alzheimer’s Disease Neuroimaging Initiative, Focused ultrasound-mediated blood-brain barrier opening in Alzheimer’s disease: long-term safety, imaging, and cognitive outcomes, J. Neurosurg. 139 (2023) 275–283. 10.3171/2022.9.JNS221565.

[26] L. Chen, E. Cruz, L.E. Oikari, P. Padmanabhan, J. Song, J. Götz, Opportunities and challenges in delivering biologics for Alzheimer’s disease by low-intensity ultrasound, Adv. Drug Deliv. Rev. 189 (2022) 114517. 10.1016/j.addr.2022.114517.

[27] J. Götz, P. Padmanabhan, Ultrasound and antibodies - a potentially powerful combination for Alzheimer disease therapy, Nat. Rev. Neurol. 20 (2024) 257–258. 10.1038/s41582-024-00943-1.

[28] Z. Fan, D. Chen, C.X. Deng, Improving ultrasound gene transfection efficiency by controlling ultrasound excitation of microbubbles, J. Control. Release Off. J. Control. Release Soc. 170 (2013) 401–413. 10.1016/j.jconrel.2013.05.039.

[29] B. Helfield, X. Chen, S.C. Watkins, F.S. Villanueva, Biophysical insight into mechanisms of sonoporation, Proc. Natl. Acad. Sci. U. S. A. 113 (2016) 9983–9988. 10.1073/pnas.1606915113.

[30] S. Roovers, G. Lajoinie, I. De Cock, T. Brans, H. Dewitte, K. Braeckmans, M. Versuis, S.C. De Smedt, I. Lentacker, Sonoprinting of nanoparticle-loaded microbubbles: Unraveling the multi-timescale mechanism, Biomaterials 217 (2019) 119250. 10.1016/j.biomaterials.2019.119250.

[31] A. van Wamel, K. Kooiman, M. Harteveld, M. Emmer, F.J. ten Cate, M. Versluis, N. de Jong, Vibrating microbubbles poking individual cells: drug transfer into cells via sonoporation, J. Control. Release Off. J. Control. Release Soc. 112 (2006) 149–155. 10.1016/j.jconrel.2006.02.007.

[32] I. Beekers, F. Mastik, R. Beurskens, P.Y. Tang, M. Vegter, A.F.W. van der Steen, N. de Jong, M.D. Verweij, K. Kooiman, High-resolution imaging of intracellular calcium fluctuations caused by oscillating microbubbles, Ultrasound Med. Biol. 46 (2020) 2017–2029. 10.1016/j.ultrasmedbio.2020.03.029.

[33] Z. Fan, R.E. Kumon, J. Park, C.X. Deng, Intracellular delivery and calcium transients generated in sonoporation facilitated by microbubbles, J. Control. Release Off. J. Control. Release Soc. 142 (2010) 31–39. 10.1016/j.jconrel.2009.09.031.

[34] Z. Fan, H. Liu, M. Mayer, C.X. Deng, Spatiotemporally controlled single cell sonoporation, Proc. Natl. Acad. Sci. U. S. A. 109 (2012) 16486–16491. 10.1073/pnas.1208198109.

[35] L.J.M. Juffermans, A. van Dijk, C.A.M. Jongenelen, B. Drukarch, A. Reijerkerk, H.E. de Vries, O. Kamp, R.J.P. Musters, Ultrasound and microbubble-induced intra- and intercellular bioeffects in primary endothelial cells, Ultrasound Med. Biol. 35 (2009) 1917–1927. 10.1016/j.ultrasmedbio.2009.06.1091.

[36] J. Park, Z. Fan, R.E. Kumon, M.E.H. El-Sayed, C.X. Deng, Modulation of intracellular Ca2+ concentration in brain microvascular endothelial cells in vitro by acoustic cavitation, Ultrasound Med. Biol. 36 (2010) 1176–1187. 10.1016/j.ultrasmedbio.2010.04.006.

[37] J. Shi, T. Han, A.C.H. Yu, P. Qin, Faster calcium recovery and membrane resealing in repeated sonoporation for delivery improvement, J. Control. Release Off. J. Control. Release Soc. 352 (2022) 385–398. 10.1016/j.jconrel.2022.10.027.

[38] J. Shi, Y. Ma, R. Shi, A.C.H. Yu, P. Qin, Manipulating long-term fates of sonoporated cells by regulating intracellular calcium for improving sonoporation-based delivery, J. Control. Release Off. J. Control. Release Soc. (2024) S0168–3659(24)00598–4. 10.1016/j.jconrel.2024.08.048.

[39] C. Jia, L. Xu, T. Han, P. Cai, A.C.H. Yu, P. Qin, Generation of reactive oxygen species in heterogeneously sonoporated cells by microbubbles with single-pulse ultrasound, Ultrasound Med. Biol. 44 (2018) 1074–1085. 10.1016/j.ultrasmedbio.2018.01.006.

[40] W. Zhong, X. Chen, P. Jiang, J.M.F. Wan, P. Qin, A.C.H. Yu, Induction of endoplasmic reticulum stress by sonoporation: linkage to mitochondria-mediated apoptosis initiation, Ultrasound Med. Biol. 39 (2013) 2382–2392. 10.1016/j.ultrasmedbio.2013.08.005.

[41] X. Chen, R.S. Leow, Y. Hu, J.M.F. Wan, A.C.H. Yu, Single-site sonoporation disrupts actin cytoskeleton organization, J. R. Soc. Interface 11 (2014) 20140071. 10.1098/rsif.2014.0071.

[42] C. Jia, J. Shi, T. Han, A.C.H. Yu, P. Qin, Spatiotemporal dynamics and mechanisms of actin cytoskeletal re-modeling in cells perforated by ultrasound-driven microbubbles, Ultrasound Med. Biol. 48 (2022) 760–777. 10.1016/j.ultrasmedbio.2021.12.014.

[43] B. Meijlink, H.R. van der Kooij, Y. Wang, H. Li, S. Huveneers, K. Kooiman, Ultrasound-activated microbubbles mediate F-actin disruptions and endothelial gap formation during sonoporation, J. Control. Release Off. J. Control. Release Soc. (2024) S0168–3659(24)00739–9. 10.1016/j.jconrel.2024.10.066.

[44] M. Wang, Y. Zhang, C. Cai, J. Tu, X. Guo, D. Zhang, Sonoporation-induced cell membrane permeabilization and cytoskeleton disassembly at varied acoustic and microbubble-cell parameters, Sci. Rep. 8 (2018) 3885. 10.1038/s41598-018-22056-8.

[45] C. Jia, J. Shi, null Yao Yao, T. Han, A.C.H. Yu, P. Qin, Plasma membrane blebbing dynamics Involved in the reversibly perforated cell by ultrasound-driven microbubbles, Ultrasound Med. Biol. 47 (2021) 733–750. 10.1016/j.ultrasmedbio.2020.11.029.

[46] R.S. Leow, J.M.F. Wan, A.C.H. Yu, Membrane blebbing as a recovery manoeuvre in site-specific sonoporation mediated by targeted microbubbles, J. R. Soc. Interface 12 (2015) 20150029. 10.1098/rsif.2015.0029.

[47] J.-Z. Zhang, J.K. Saggar, Z.-L. Zhou, B. Hu, Different effects of sonoporation on cell morphology and viability, Bosn. J. Basic Med. Sci. 12 (2012) 64–68. 10.17305/bjbms.2012.2497.

[48] B. Helfield, X. Chen, S.C. Watkins, F.S. Villanueva, Transendothelial perforations and the sphere of influence of single-site sonoporation, Ultrasound Med. Biol. 46 (2020) 1686–1697. 10.1016/j.ultrasmedbio.2020.02.017.

[49] I. Beekers, S.A.G. Langeveld, B. Meijlink, A.F.W. van der Steen, N. de Jong, M.D. Verweij, K. Kooiman, Internalization of targeted microbubbles by endothelial cells and drug delivery by pores and tunnels, J. Control. Release Off. J. Control. Release Soc. 347 (2022) 460–475. 10.1016/j.jconrel.2022.05.008.

[50] A. Abrahao, Y. Meng, M. Llinas, Y. Huang, C. Hamani, T. Mainprize, I. Aubert, C. Heyn, S.E. Black, K. Hynynen, N. Lipsman, L. Zinman, First-in-human trial of blood-brain barrier opening in amyotrophic lateral sclerosis using MR-guided focused ultrasound, Nat. Commun. 10 (2019) 4373. 10.1038/s41467-019-12426-9.

[51] Y. Meng, C.B. Pople, H. Lea-Banks, A. Abrahao, B. Davidson, S. Suppiah, L.M. Vecchio, N. Samuel, F. Mahmud, K. Hynynen, C. Hamani, N. Lipsman, Safety and efficacy of focused ultrasound induced blood-brain barrier opening, an integrative review of animal and human studies, J. Control. Release Off. J. Control. Release Soc. 309 (2019) 25–36. 10.1016/j.jconrel.2019.07.023.

[52] Y. Huang, R. Alkins, M.L. Schwartz, K. Hynynen, Opening the blood-brain barrier with MR imaging-guided focused ultrasound: preclinical testing on a trans-human skull porcine model, Radiology 282 (2017) 123–130. 10.1148/radiol.2016152154.

[53] T. Ilovitsh, A. Ilovitsh, J. Foiret, C.F. Caskey, J. Kusunose, B.Z. Fite, H. Zhang, L.M. Mahakian, S. Tam, K. Butts-Pauly, S. Qin, K.W. Ferrara, Enhanced microbubble contrast agent oscillation following 250 kHz insonation, Sci. Rep. 8 (2018) 16347. 10.1038/s41598-018-34494-5.

[54] T. Ilovitsh, Y. Feng, J. Foiret, A. Kheirolomoom, H. Zhang, E.S. Ingham, A. Ilovitsh, S.K. Tumbale, B.Z. Fite, B. Wu, M.N. Raie, N. Zhang, A.J. Kare, M. Chavez, L.S. Qi, G. Pelled, D. Gazit, O. Vermesh, I. Steinberg, S.S. Gambhir, K.W. Ferrara, Low-frequency ultrasound-mediated cytokine transfection enhances T cell recruitment at local and distant tumor sites, Proc. Natl. Acad. Sci. U. S. A. 117 (2020) 12674–12685. 10.1073/pnas.1914906117.

[55] I. Beekers, M. Vegter, K.R. Lattwein, F. Mastik, R. Beurskens, A.F.W. van der Steen, N. de Jong, M.D. Verweij, K. Kooiman, Opening of endothelial cell-cell contacts due to sonoporation, J. Control. Release Off. J. Control. Release Soc. 322 (2020) 426–438. 10.1016/j.jconrel.2020.03.038.

[56] B.R. Brückner, A. Janshoff, Importance of integrity of cell-cell junctions for the mechanics of confluent MDCK II cells, Sci. Rep. 8 (2018) 14117. 10.1038/s41598-018-32421-2.

[57] L. Chen, R. Sutharsan, J.L. Lee, E. Cruz, B. Asnicar, T. Palliyaguru, J.M. Wasielewska, A. Gaudin, J. Song, G. Leinenga, J. Götz, Claudin-5 binder enhances focused ultrasound-mediated opening in an in vitro blood-brain barrier model, Theranostics 12 (2022) 1952–1970. 10.7150/thno.65539.

[58] L. Chen, J. Song, G. Richter-Stretton, W. Lee, P. Padmanabhan, J. Götz, Multimodal evaluation of blood-brain barrier opening in mice in response to low-intensity ultrasound and a claudin-5 binder, Nanotheranostics 8 (2024) 427–441. 10.7150/ntno.95146.

[59] P.S. Caceres, D. Gravotta, P.J. Zager, N. Dephoure, E. Rodriguez-Boulan, Quantitative proteomics of MDCK cells identify unrecognized roles of clathrin adaptor AP-1 in polarized distribution of surface proteins, Proc. Natl. Acad. Sci. U. S. A. 116 (2019) 11796–11805. 10.1073/pnas.1821076116.

[60] Y.-S. Chen, R.A. Mathias, S. Mathivanan, E.A. Kapp, R.L. Moritz, H.-J. Zhu, R.J. Simpson, Proteomics profiling of Madin-Darby canine kidney plasma membranes reveals Wnt-5a involvement during oncogenic H-Ras/TGF-beta-mediated epithelial-mesenchymal transition, Mol. Cell. Proteomics MCP 10 (2011) M110.001131. 10.1074/mcp.M110.001131.

[61] T.-W. Chen, T.J. Wardill, Y. Sun, S.R. Pulver, S.L. Renninger, A. Baohan, E.R. Schreiter, R.A. Kerr, M.B. Orger, V. Jayaraman, L.L. Looger, K. Svoboda, D.S. Kim, Ultrasensitive fluorescent proteins for imaging neuronal activity, Nature 499 (2013) 295–300. 10.1038/nature12354.

[62] D. Andrade, J. Rosenblatt, Apoptotic regulation of epithelial cellular extrusion, Apoptosis Int. J. Program. Cell Death 16 (2011) 491–501. 10.1007/s10495-011-0587-z.

[63] K. Duszyc, J.B. von Pein, D. Ramnath, D. Currin-Ross, S. Verma, F. Lim, M.J. Sweet, K. Schroder, A.S. Yap, Apical extrusion prevents apoptosis from activating an acute inflammatory program in epithelia, Dev. Cell 58 (2023) 2235–2248.e6. 10.1016/j.devcel.2023.08.009.

[64] S.M. Stamatovic, R.F. Keep, M.M. Wang, I. Jankovic, A.V. Andjelkovic, Caveolae-mediated internalization of occludin and claudin-5 during CCL2-induced tight junction remodeling in brain endothelial cells, J. Biol. Chem. 284 (2009) 19053–19066. 10.1074/jbc.M109.000521.

[65] D. Kamin, M.A. Lauterbach, V. Westphal, J. Keller, A. Schönle, S.W. Hell, S.O. Rizzoli, High- and low-mobility stages in the synaptic vesicle cycle, Biophys. J. 99 (2010) 675–684. 10.1016/j.bpj.2010.04.054.

[66] P. Padmanabhan, A. Kneynsberg, E. Cruz, R. Amor, J.-B. Sibarita, J. Götz, Single-molecule imaging reveals Tau trapping at nanometer-sized dynamic hot spots near the plasma membrane that persists after microtubule perturbation and cholesterol depletion, EMBO J. 41 (2022) e111265. 10.15252/embj.2022111265.

[67] V. Westphal, S.O. Rizzoli, M.A. Lauterbach, D. Kamin, R. Jahn, S.W. Hell, Video-rate far-field optical nanoscopy dissects synaptic vesicle movement, Science 320 (2008) 246–249. 10.1126/science.1154228.

[68] R.E. Kumon, M. Aehle, D. Sabens, P. Parikh, D. Kourennyi, C.X. Deng, Ultrasound-induced calcium oscillations and waves in Chinese hamster ovary cells in the presence of microbubbles, Biophys. J. 93 (2007) L29–31. 10.1529/biophysj.107.113365.

[69] H.V. Goodson, E.M. Jonasson, Microtubules and microtubule-associated proteins, Cold Spring Harb. Perspect. Biol. 10 (2018) a022608. 10.1101/cshperspect.a022608.

[70] R.A. Larsen, C. Cusumano, A. Fujioka, G. Lim-Fong, P. Patterson, J. Pogliano, Treadmilling of a prokaryotic tubulin-like protein, TubZ, required for plasmid stability in Bacillus thuringiensis, Genes Dev. 21 (2007) 1340–1352. 10.1101/gad.1546107.

[71] P. Fan, Y. Zhang, X. Guo, C. Cai, M. Wang, D. Yang, Y. Li, J. Tu, L.A. Crum, J. Wu, D. Zhang, Cell-cycle-specific cellular responses to sonoporation, Theranostics 7 (2017) 4894–4908. 10.7150/thno.20820.

[72] D.G. Blackmore, D. Razansky, J. Götz, Ultrasound as a versatile tool for short- and long-term improvement and monitoring of brain function, Neuron 111 (2023) 1174–1190. 10.1016/j.neuron.2023.02.018.

[73] R. Haugse, A. Langer, E.T. Murvold, D.E. Costea, B.T. Gjertsen, O.H. Gilja, S. Kotopoulis, G. Ruiz de Garibay, E. McCormack, Low-intensity sonoporation-induced intracellular signalling of pancreatic cancer cells, fibroblasts and endothelial cells, Pharmaceutics 12 (2020) 1058. 10.3390/pharmaceutics12111058.

[74] T. van Rooij, I. Skachkov, I. Beekers, K.R. Lattwein, J.D. Voorneveld, T.J.A. Kokhuis, D. Bera, Y. Luan, A.F.W. van der Steen, N. de Jong, K. Kooiman, Viability of endothelial cells after ultrasound-mediated sonoporation: Influence of targeting, oscillation, and displacement of microbubbles, J. Control. Release Off. J. Control. Release Soc. 238 (2016) 197–211. 10.1016/j.jconrel.2016.07.037.

[75] K. Kooiman, M. Foppen-Harteveld, A.F.W. van der Steen, N. de Jong, Sonoporation of endothelial cells by vibrating targeted microbubbles, J. Control. Release Off. J. Control. Release Soc. 154 (2011) 35–41. 10.1016/j.jconrel.2011.04.008.

[76] M. Cattaneo, G. Guerriero, G. Shakya, L.A. Krattiger, L. G. Paganella, M.L. Narciso, O. Supponen, Cyclic jetting enables microbubble-mediated drug delivery, Nat. Phys. (2025). 10.1038/s41567-025-02785-0.

[77] W. Marsh, Propidium, in: XPharm Compr. Pharmacol. Ref., Elsevier, 2007: pp. 1–2. 10.1016/B978-008055232-3.62478-X.

[78] J. Huff, A. Bergter, B. Luebbers, Multiplex mode for the LSM 9 series with Airyscan 2: fast and gentle confocal super-resolution in large volumes, Nat. Methods (2019).

[79] X. Wu, J.A. Hammer, ZEISS Airyscan: optimizing usage for fast, gentle, super-resolution imaging, Methods Mol. Biol. Clifton NJ 2304 (2021) 111–130. 10.1007/978-1-0716-1402-0_5.

[80] J.-Y. Tinevez, N. Perry, J. Schindelin, G.M. Hoopes, G.D. Reynolds, E. Laplantine, S.Y. Bednarek, S.L. Shorte, K.W. Eliceiri, TrackMate: An open and extensible platform for single-particle tracking, Methods San Diego Calif 115 (2017) 80–90. 10.1016/j.ymeth.2016.09.016.

[81] P. Padmanabhan, A. Kneynsberg, E. Cruz, A. Briner, J. Götz, Single-molecule imaging of Tau reveals how phosphorylation affects its movement and confinement in living cells, Mol. Brain 17 (2024) 7. 10.1186/s13041-024-01078-6.

[82] P. Padmanabhan, R. Martínez-Mármol, D. Xia, J. Götz, F.A. Meunier, Frontotemporal dementia mutant Tau promotes aberrant Fyn nanoclustering in hippocampal dendritic spines, eLife 8 (2019) e45040. 10.7554/eLife.45040.

