## Supplementary Material for "High-resolution imaging reveals a cascade of interconnected cellular bioeffects differentiating the long-term fates of sonoporated cells"

**[*]Correspondence:**

### SUPPLEMENTARY VIDEO LEGEND

**Video S1. Changes in intracellular calcium and propidium iodide in a non-sonoporated cell (related to Figure 4).** Time-lapse video of GCaMP6f (left) and PI (right) signals in a representative non-sonoporated GCaMP6f-MDCK II cell. Time is shown in min:s. The display rate is 30 frames per second.

**Video S2. Changes in intracellular calcium and propidium iodide in a reversibly sonoporated cell (related to Figure 4).** Time-lapse video of GCaMP6f (left) and PI (right) signals in a representative reversibly sonoporated GCaMP6f-MDCK II cell. Time is shown in min:s. The display rate is 30 frames per second.

**Video S3. Changes in intracellular calcium and propidium iodide in an irreversibly sonoporated cell (related to Figure 4).** Time-lapse video of GCaMP6f (left) and PI (right) signals in a representative irreversibly sonoporated GCaMP6f-MDCK II cell. Time is shown in min:s. The display rate is 30 frames per second.

**Video S4. Changes in cell size, CLDN5-containing vesicle dynamics, and propidium iodide signal in a non-sonoporated cell (related to Figure 5).** Time-lapse video of EGFP-CLDN5 (left) and PI (right) signals in a representative non-sonoporated EGFP-CLDN5-MDCK II cell. Time is shown in min:s. The display rate is 30 frames per second.

**Video S5. Changes in cell size, CLDN5-containing vesicle dynamics, and propidium iodide signal in a reversibly sonoporated cell (related to Figure 5).** Time-lapse video of EGFP-CLDN5 (left) and PI (right) signals in a representative reversibly sonoporated EGFP-CLDN5-MDCK II cell. Time is shown in min:s. The display rate is 30 frames per second.

**Video S6. Changes in cell size, CLDN5-containing vesicle dynamics, and propidium iodide signal in an irreversibly sonoporated cell (related to Figure 5).** Time-lapse video of EGFP-CLDN5 (left) and PI (right) signals in a representative irreversibly sonoporated EGFP-CLDN5-MDCK II cell. Time is shown in min:s. The display rate is 30 frames per second.

**Video S7. Changes in cell size, microtubule network dynamics and propidium iodide signal in a non-sonoporated cell (related to Figure 6).** Time-lapse video of EGFP-CLDN5 (left), SiR-Tubulin (middle) and PI (right) signals in a representative non-sonoporated EGFP-CLDN5 MDCK II cell. The microtubule network was stained with SiR-Tubulin. Time is shown in min:s. The display rate is 30 frames per second.

**Video S8. Changes in cell size, microtubule network dynamics and propidium iodide signal in a reversibly sonoporated cell (related to Figures 6 and 7).** Time-lapse video of EGFP-CLDN5 (left), SiR-Tubulin (middle) and PI (right) signals in a representative reversibly sonoporated EGFP-CLDN5 MDCK II cell. The microtubule network was stained with SiR-Tubulin. Time is shown in min:s. The display rate is 30 frames per second.

**Video S9. Changes in cell size, microtubule network dynamics and propidium iodide signal in an irreversibly sonoporated cell (related to Figure 6).** Time-lapse video of EGFP-CLDN5 (left), SiR-Tubulin (middle) and PI (right) signals in a representative irreversibly sonoporated EGFP-CLDN5 MDCK II cell. The microtubule network was stained with SiR-Tubulin. Time is shown in min:s. The display rate is 30 frames per second.

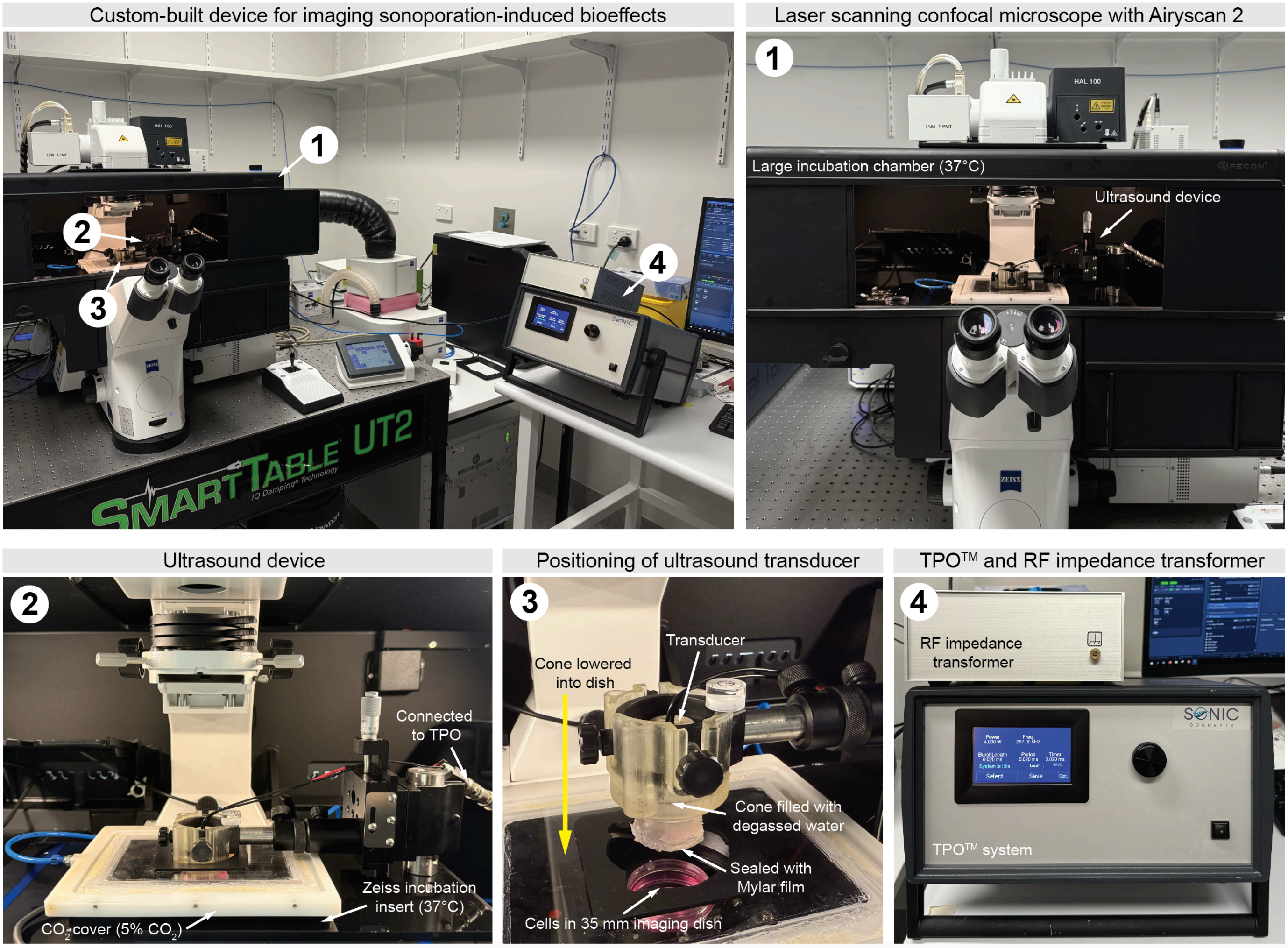


**Supplementary Figure 1. Custom-built high-resolution imaging device for investigating bioeffects triggered by ultrasound-induced sonoporation.** A collection of images of the entire setup comprising of the laser scanning confocal microscope with non-linear optics and Airyscan 2 detector, a single-element ultrasound transducer with coupling cone, temperature, $CO_2$ and air humidifier controls, and the TPO system.

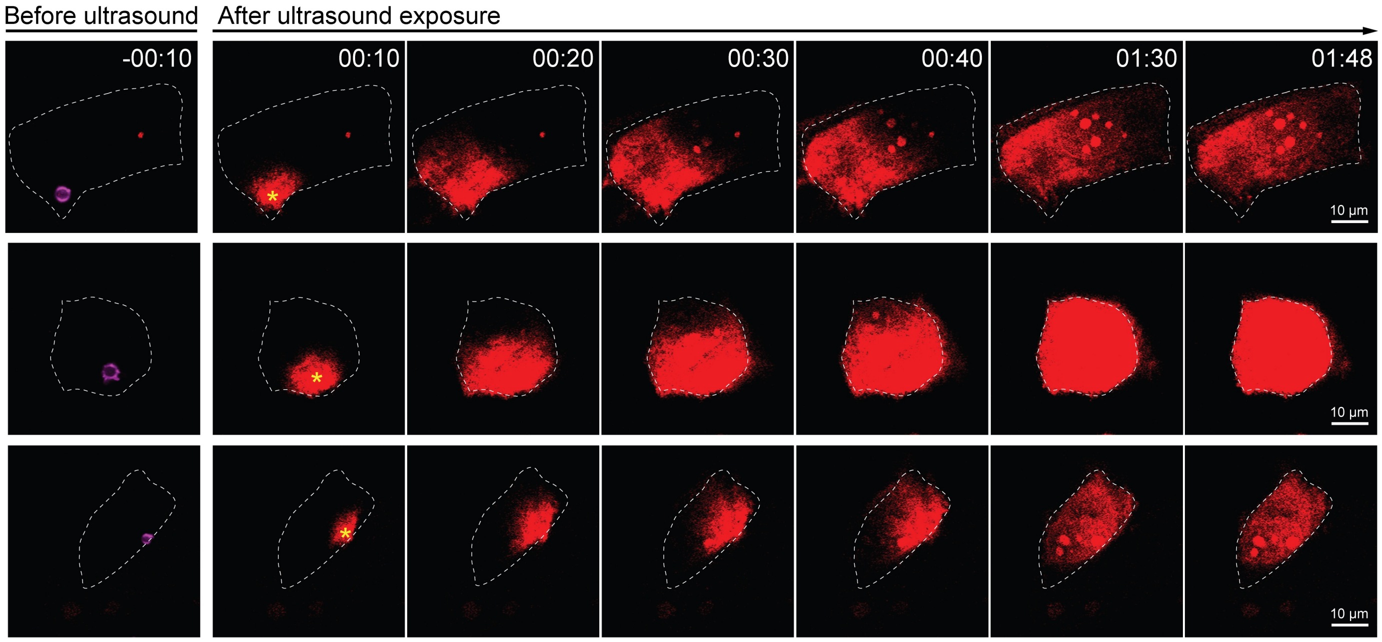


**Supplementary Figure 2. The site of PI uptake coincides with the location of the CD51-targeted microbubble.** Representative time-series images of PI uptake in three GCaMP6f-MDCK II cells. The z-plane above the cell where the CD51-targeted microbubble was clearly visible was captured before sonication and overlayed on the PI images before sonication to indicate the microbubble position. The yellow asterisk in the first frame after sonication shows the microbubble location determined before sonication. Time is shown in min:s.

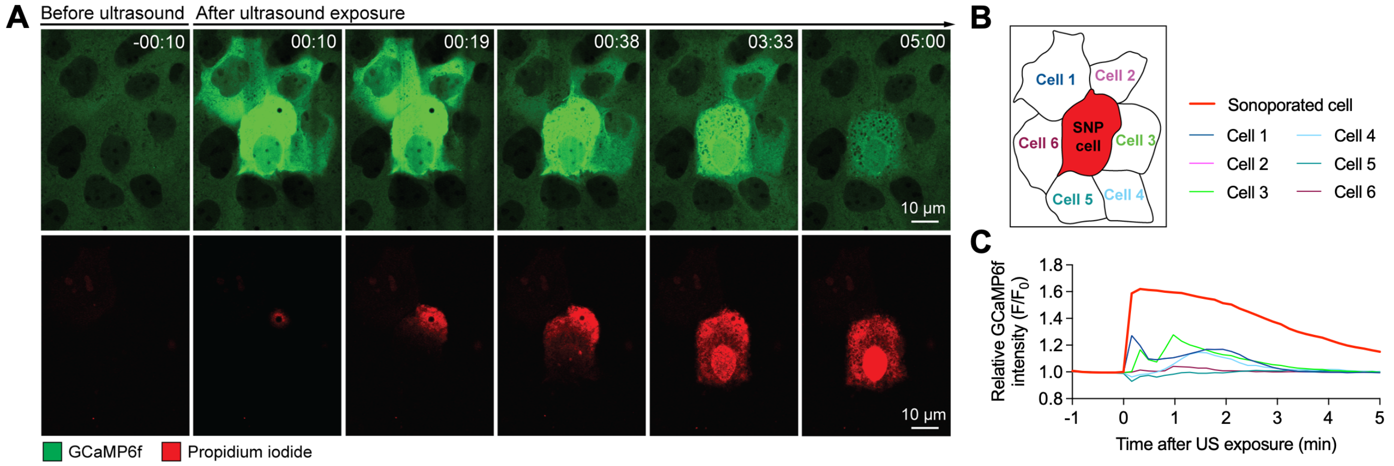


**Supplementary Figure 3. The spreading of calcium waves to cells adjacent to a sonoporated cell. (A)** Time-lapse images of GCaMP6f fluorescence intensity representing intracellular calcium level (top panels) and the corresponding PI level (bottom panels) before and after sonication in a sonoporated cell and its adjacent non-sonoporated cells. **(B-C)** The outline (B) and the relative GCaMP6f intensity (C) of the sonoporated cell in A and its six adjacent non-sonoporated cells.
